# Rac1 signaling in microglia is essential for synaptic proteome plasticity and experience-dependent cognitive performance

**DOI:** 10.1101/2021.10.18.464802

**Authors:** Renato Socodato, Tiago O. Almeida, Camila C. Portugal, Evelyn C. S. Santos, Joana Tedim-Moreira, Teresa Canedo, Filipa I. Baptista, Ana Magalhães, António F. Ambrósio, Cord Brakebusch, Boris Rubinstein, Irina S. Moreira, Teresa Summavielle, Inês Mendes Pinto, João B. Relvas

**Affiliations:** Institute of Research and Innovation in Health (i3S) and Institute for Molecular and Cell Biology (IBMC), University of Porto, Porto, Portugal; ESS.PP, Escola Superior de Saúde do Politécnico do Porto, Porto, Portugal; International Iberian Nanotechnology Laboratory (INL), Braga, Portugal; Center for Innovative Biomedicine and Biotechnology (CIBB), Coimbra Institute for Clinical and Biomedical Research (iCBR), and Clinical Academic Center of Coimbra (CACC), University of Coimbra, Coimbra, Portugal; Faculty of Medicine of the University of Porto (FMUP), Porto, Portugal; Stowers Institute for Medical Research; Kansas City, MO, USA; Department of Life Sciences, Center for Innovative Biomedicine and Biotechnology (CIBB) and CNC-Center for Neuroscience and Cell Biology, University of Coimbra, Coimbra, Portugal; Molecular Pathology Section, BRIC, Københavns Biocenter, Copenhagen, Denmark; ICBAS - School of Medicine and Biomedical Sciences, Porto, Portugal

## Abstract

Microglial homeostatic functions are fundamental to regulate the central nervous system microenvironment. We use conditional cell-specific gene targeting, RNA-seq profiling, high-throughput proteomics, phosphoproteomics, systems biology, and animal behavior to report a critical role for the RhoGTPase Rac1 in regulating adult microglia physiology. Ablation of Rac1 in adult microglia impaired their ability to sense and interpret the brain microenvironment and affected their capacity to communicate with synapses to drive cognitive performance, both at the steady-state and during experience-dependent plasticity. Overall, our results reveal a novel and central role for Rac1 as a regulator of microglia homeostasis and a molecular driver of the microglia-synapse crosstalk required for context-dependent sociability and learning related to memory.

## Introduction

Microglia, the largest myeloid cell population in the CNS, are best known for their protective roles in the brain. Not only are microglia involved in physiological synaptic pruning, but they also secrete a plethora of chemical mediators (including neurotransmitters and growth factors) that affects brain synapses ^1^. Targeted gene ablation experiments indicate that microglia-secreted factors are putative modulators of synaptic activity and plasticity ^2-5^. Even though the microglial secretome conceivably regulates many aspects of microglia-synapse crosstalk, little is known about the magnitude and specificity of the alterations in synaptic signaling and remodeling elicited by the chemical mediators secreted by microglia ^6^.

Remodeling of the synaptic structure and signaling network, bona fide synaptic plasticity modalities, occurs continuously throughout life and underlies high-order cognitive functions necessary for an organism to adapt to environmental inputs ^7^. Thus, deciphering how the brain encodes and stores experiences triggered by different environmental contexts requires in-depth knowledge of the cellular and molecular mechanisms governing synaptic remodeling, including those in which microglia participate.

The GTPases of the Rho family, including the most well-characterized members Rac1, RhoA, and Cdc42, are critical orchestrators of process extension required for cell-cell communication and intracellular trafficking involved in secretion of chemical mediators ^8^, making them likely players to govern microglial homeostatic functions. Studies with myeloid cells from outside the CNS have identified a central role for Rac signaling in executing the inflammatory program of the innate immune system ^9^. There are three Rac isoforms, but only Rac1 and 2 are significantly expressed in myeloid cells ^10^. Gene-targeting approaches revealed essential modulatory roles for Rac1 in the CNS, including axonal growth and stability ^11^, oligodendrocyte myelination ^12^, and astrocyte morphology ^13^. Yet, no information is currently available on Rac1 regulation of microglial homeostatic functions in vivo.

Here, by combining conditional cell-specific gene ablation, RNA-seq profiling, high-throughput proteomics, single-cell live imaging, systems biology, and animal behavior, we revealed a multiscale role for microglial Rac1 in the adult brain. Ablation of Rac1 specifically suppressed the ability of microglia to sense the brain microenvironment, severing the capacity of microglia to communicate with synapses properly. Loss of homeostatic microglia-to-synapse communication primarily disrupted the remodeling of the proteome network of synapses required for experience-dependent plasticity, impairing context-dependent learning, memory, and social interaction.

## Results

### Conditional ablation of Rac1 in microglia

We have reported that the RhoGTPase RhoA is a critical negative regulator of microglia immune activity causing neurodegeneration ^4^. Here, we sought to demonstrate that targeting Rac1 disrupts microglia homeostasis and consequently microglia-to-synaptic communication related to cognitive performance. Thus, we crossed Cx3cr1^CreER-IRES-EYFP^ mice ^3, 14, 15^ with mice carrying Rac1 floxed alleles ^16^ (**Suppl. Fig. 1A**). In Cx3cr1^CreER+^ mice, the CreER-IRES-EYFP transgene is transcriptionally active in Iba1^+^ brain myeloid cells ^17^ and microglia ^3, 4^. Following tamoxifen administration, Cre migrates to the nucleus inducing Rac1 gene inactivation in Cx3cr1^CreER+^:Rac1^fl/fl^ mice.

**Figure 1.**
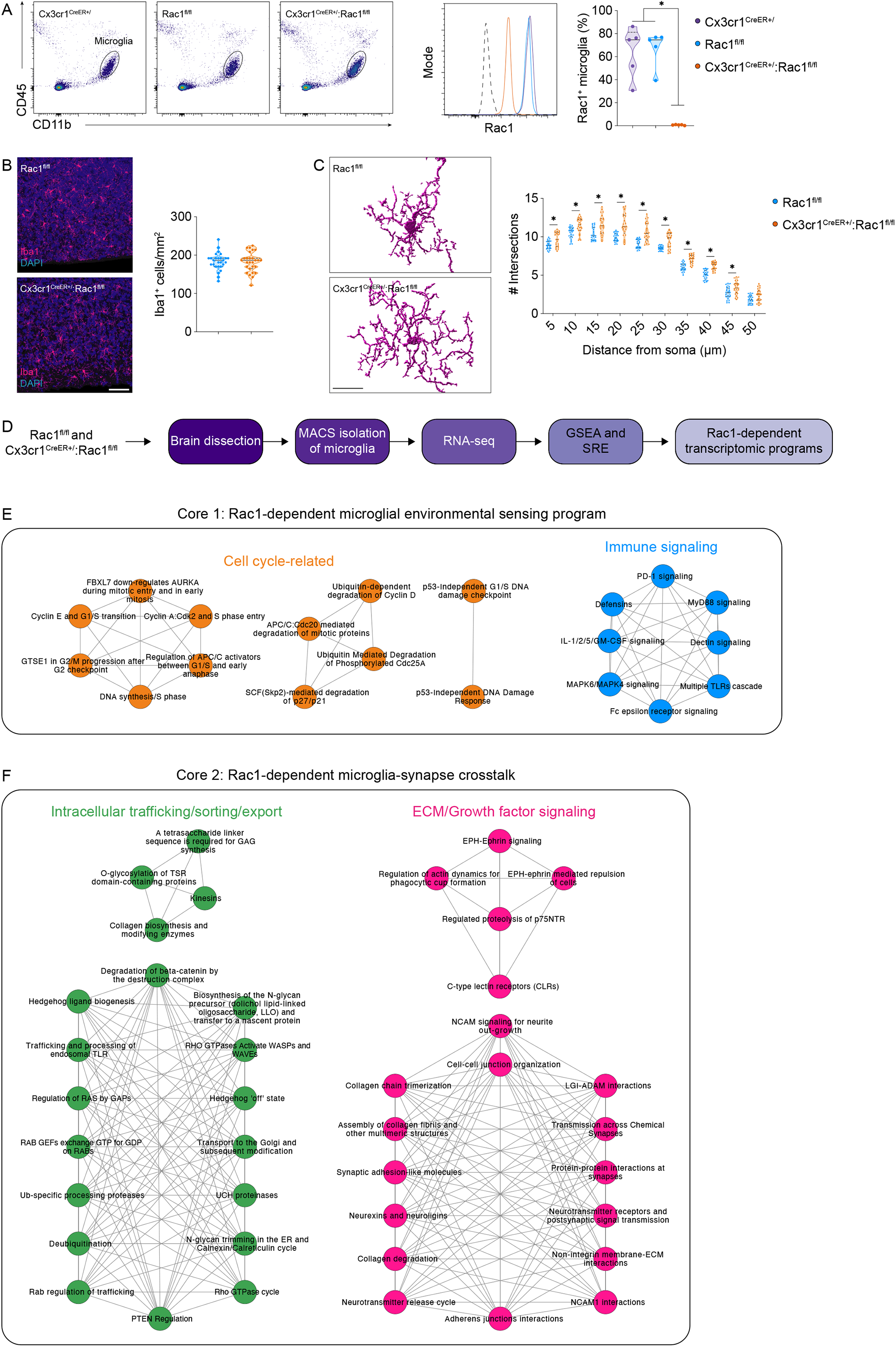
The microglial Rac1 signature. **A**, Flow cytometry analysis of Rac1 expression in microglia from Cx3cr1^CreER+^, Rac1^fl/fl^, and Cx3cr1^CreER+^:Rac1^fl/fl^ mice (n=5 animals per genotype). Violin plots (median and quartiles) depict Rac1^+^ microglia. *P<0.05 (One-way ANOVA). **B and C**, Histological confocal analysis and Imaris rendering of Iba1 on tissue sections from neocortices of Rac1^fl/fl^ and Cx3cr1^CreER+^:Rac1^fl/fl^ mice (n= 3 slices from 9 animals per genotype (B) or n=25 cells from 3 different mice per genotype (C)). Violin plots (median and quartiles) depict Iba1^+^ cells (B) or Sholl analysis (C). *P<0.05 (Two-way ANOVA comparing each distance between genotypes). Scale bars: B: 50 μm; C: 10 μm. **D**, Workflow of microglial isolation, sequencing, and downstream bioinformatics analysis. **E and F**, Core Rac1-dependent microglia transcriptomic programs revealed by analogy concatenation of GSEA and SRA (n=3 mice per genotype). Interaction networks display the top enriched pathways within each module.

To investigate the function of Rac1 in adult microglia, we administered tamoxifen by oral gavage to 4-5 weeks-old control (Rac1^fl/fl^ or Cx3cr1^CreER+^) and conditional microglia Rac1 (Cx3cr1^CreER+^:Rac1^fl/fl^) mice and analyzed their brains 6-8 weeks later. While Cx3cr1^CreER+^ mice have only one intact copy of the Cx3cr1 allele, haploinsufficiency had no impact on Rac1 expression in Cx3cr1^CreER+^ microglia (**Fig. 1A**) and causes no overt effects on microglial reactivity, synapse number, or memory ^4^. Analysis by flow cytometry (gating strategy in **Suppl. Fig. 1B**) confirmed a substantial decrease of Rac1 protein amounts in Cx3cr1^CreER+^:Rac1^fl/fl^ microglia compared to Rac1^fl/fl^ or Cx3cr1^CreER+^ microglia (**Fig. 1A**). We also corroborated the loss of Rac1 in Cx3cr1^CreER+^:Rac1^fl/fl^ microglia by qRT-PCR in flow cytometry-sorted microglia (**Suppl. Fig. 1C**).

### Profiling Rac1-dependent changes in microglia

We then evaluated the requirement of Rac1 for microglia homeostasis in the adult brain. Using Iba1 immunocytochemistry, we observed no difference in the number of Iba1^+^ microglia between the brains of Cx3cr1^CreER+^:Rac1^fl/fl^ and Rac1^fl/fl^ mice (**Fig. 1B**). Morphology changes are associated with alterations in microglial cell function, and morphological analysis of brain tissue sections using Iba1 revealed that Cx3cr1^CreER+^:Rac1^fl/fl^ microglia were hyper-ramified compared with microglia from Rac1^fl/fl^ littermates (**Fig. 1C**).

To further investigate how the ablation of Rac1 affects microglia, we conducted RNA-seq analysis in MACS-separated microglia from the brains of Cx3cr1^CreER+^:Rac1^fl/fl^ and Rac1^fl/fl^ littermates (**Fig. 1D**). Contingency analyses (**Suppl. Fig. 1D**) using known microglial signature modules (including homeostatic ^18^, pro-inflammatory ^19^, oxidative stress ^20^, disease-associated ^21^, injury-related ^22^, and aging ^22^) showed no significant association with the Rac1-regulated differentially expressed genes (DEGs) (**Suppl. Table 1)**. To pinpoint the most relevant biological pathways altered in the microglial transcriptome after Rac1 ablation, we performed pre-ranked gene set enriched analysis (GSEA) and String Ranked Enrichment (SRE) (**Fig. 1D**). GSEA and SRE revealed significant upregulation of 57 and downregulation of 198 biological pathways in microglia upon Rac1 ablation (**Suppl. Table 2**). Manual trimming and analogy recategorizing relevant pathways to build logical signaling modalities led to the classification of two major microglial transcriptomic programs regulated by Rac1: microglial environmental sensing (**Fig. 1E**) and microglia-synapse crosstalk (**Fig. 1F**).

### Rac1 is necessary for microglial environmental sensing

The first core transcriptomic program — microglial environmental sensing — included two modules: cell cycle regulation and innate immune signaling (**Fig. 1E** and **Suppl. Table 3**). The cell cycle-related module included pathways critical for cell cycle progression and fidelity (**Fig. 1E** and **Suppl. Table 3**). The innate immune signaling module included pathways leading to MAPK and NFkB activation, transduction of interleukin 1 and 2 signaling cascades, and signaling through Toll-like receptors (**Fig. 1E** and **Suppl. Table 3**). We confirmed some of the transcripts’ altered expression related to microglia immune signaling, including Csf1r, Spi1, C1qb, Pros1, Mertk, Trem2, Tlr2, and Tlr7, by qRT-PCR in flow cytometry-sorted microglia (**Suppl. Fig. 2**). Overall, the transcriptomic data suggest that the ablation of Rac1 interferes with the microglial response to external cues.

To functionally validate that microglial response to external cues requires Rac1 signaling, we used fluorescent biosensors and live-cell imaging in microglial cultures. We first tested the effect of LPS, which produces a classical pro-inflammatory cascade in microglia. As expected, we observed increased production of reactive oxygen species (ROS; using the HSP33-cys FRET-based biosensor ^4, 23, 24^) in control (scrambled) microglia exposed to LPS (**Fig. 2A**). However, the knockdown (KD) of Rac1 (validation in **Suppl. Table 4**) significantly attenuated the LPS effect in increasing microglial ROS generation (**Fig. 2A**). In myeloid cells, ROS production may lead to NFkB activation ^25^, initiating inflammatory signaling ^26^. Accordingly, we found, using an NFkB pathway inhibitor biosensor ^4, 27^, that knocking down Rac1 significantly prevented the activation of the NFkB pathway by LPS (**Fig. 2B**). Second, we studied lipid sensing signaling in Rac1-deficient microglia. To analyze the lipid sensing cascade, we exposed microglia to phosphatidylcholine (PC), an essential lipid found in neuronal membranes, and measured, in real-time, the local production of diacylglycerol (Dag; a second messenger generated by phospholipase C activation) and the mobilization of Ca^2+^ from the endoplasmic reticulum (ER) into the cytosol. Using the DagLas biosensor ^28^, we observed that PC caused a significant increase in Dag production in the processes of control microglia, an effect abrogated in Rac1 KD microglia (**Fig. 2C**). Using the D1ER biosensor ^29^, we also found that Rac1 KD significantly blocked the PC-induced release of Ca^2+^ from the ER into the cytosol of microglia (**Fig. 2D**). Lastly, we studied ATP signaling, a classical danger-associated molecular pattern, in Rac1-deficient microglia. We monitored global cytosolic Ca^2+^ mobilization and PKC activity in living microglia to study ATP signaling. Using the Ca2+ sensor RGECO ^30^ and the FRET-based PKC activity reporter CKAR ^31^, we observed that ATP increased Ca2+ mobilization and PKC activation in microglia overexpressing wild-type Rac1 (**Fig. 2E** and **F**); however, overexpressing a dominant-negative Rac1 mutant (Rac1^T17N^) attenuated the ATP effect significantly (**Fig. 2E** and **F**).

**Figure 2.**
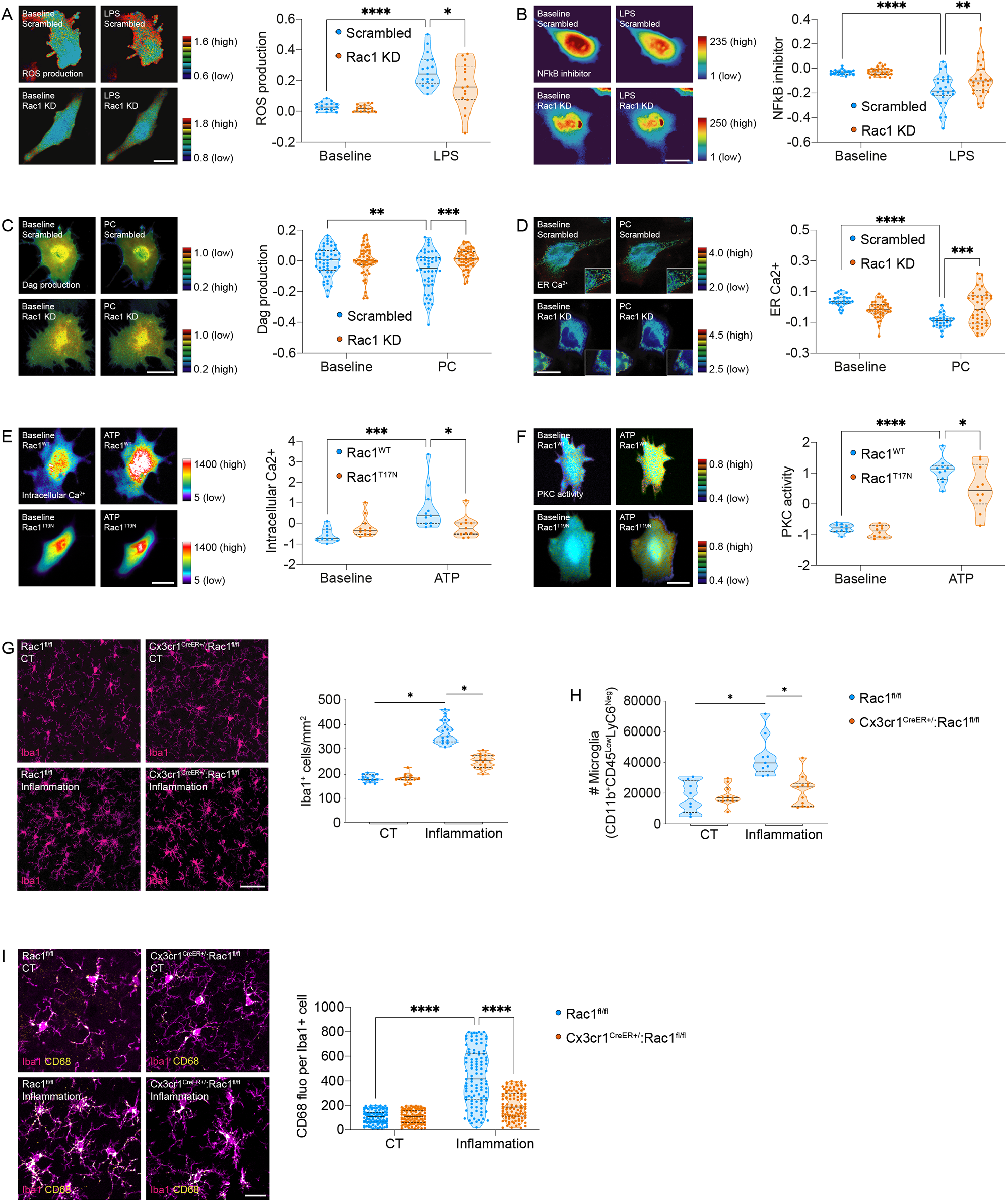
Rac1 is a relay switch for microglial sensing of external cues. **A**, Control (scrambled) or Rac1 knocked down (KD) HMC3 microglia expressing a ROS FRET biosensor were recorded in saline before (t=0 min; baseline) and after treatment with 100 ng/ml LPS. Violin plots (median and quartiles) show time-lapse Donor/FRET ratio changes (n=16-17 cells pooled across 3 different experiments). Pseudocolor ramps display representative ratio values. *p<0.05; ****p<0.0001 (Two-Way ANOVA). Scale bar: 10 μm. **B**, Control (scrambled) or Rac1 knocked down (KD) HMC3 microglia expressing an NFkB inhibitor biosensor were recorded in saline before (t=0 min; baseline) and after treatment with 100 ng/ml LPS. Violin plots (median and quartiles) show time-lapse fluorescence changes (n=25-31 cells pooled across 3 different experiments). Pseudocolor ramps display representative fluorescence values. **p<0.01; ****p<0.0001 (Two-Way ANOVA). Scale bar: 5 μm. **C**, Control (scrambled) or Rac1 knocked down (KD) HMC3 microglia expressing a diacylglycerol (Dag) FRET biosensor were recorded in saline before (t=0 min; baseline) and after treatment with 10 ng/ml phosphatidylcholine (PC). Violin plots (median and quartiles) show time-lapse FRET/Donor ratio changes (n=50-58 cells pooled across 3 different experiments). Pseudocolor ramps display representative ratio values. **p<0.01; ***p<0.001 (Two-Way ANOVA). Scale bar: 5 μm. **D**, Control (scrambled) or Rac1 knocked down (KD) HMC3 microglia expressing an endoplasmic reticulum (ER) Ca2+ FRET biosensor were recorded in saline before (t=0 min; baseline) and after treatment with 10 ng/ml PC. Violin plots (median and quartiles) show time-lapse FRET/Donor ratio changes (n=26-42 cells pooled across 3 different experiments). Pseudocolor ramps display representative ratio values. ***p<0.001; ****p<0.0001 (Two-Way ANOVA). Scale bar: 5 μm. **E**, Control (Rac1^WT^) or Rac1 dominant-negative (Rac1^T17N^) HMC3 microglia expressing a cytosolic Ca2+ biosensor were recorded in saline before (t=0 min; baseline) and after treatment with 100μM ATP. Violin plots (median and quartiles) show time-lapse fluorescence changes (n=11 cells per group pooled across 3 different experiments). Pseudocolor ramps display representative fluorescence values. *p<0.05; ***p<0.001 (Two-Way ANOVA). Scale bar: 20 μm. **F**, Control (Rac1^WT^) or Rac1 dominant-negative (Rac1^T17N^) HMC3 microglia expressing a PKC FRET biosensor were recorded in saline before (t=0 min; baseline) and after treatment with 100μM ATP. Violin plots (median and quartiles) show FRET/Donor ratio changes (n=10 cells per group pooled across 3 different experiments). Pseudocolor ramps display representative ratio values. *p<0.05; ****p<0.0001 (Two-Way ANOVA). Scale bar: 20 μm. **G**, Histological confocal analysis of Iba1 on tissue sections from neocortices of Rac1^fl/fl^ and Cx3cr1^CreER+^:Rac1^fl/fl^ mice after treatment with saline (CT) or intraperitoneal injection of 4 μg/kg LPS (inflammation) for 24 h (n=3 slices from 4 animals per group). Violin plots (median and quartiles) depict Iba1^+^ cell counts. *P<0.05 (Two-way ANOVA). Scale bar: 50 μm. **H**, Flow cytometry analysis of microglia from the brains of Rac1^fl/fl^ and Cx3cr1^CreER+^:Rac1^fl/fl^ mice after treatment with saline (CT) or intraperitoneal injection of 4 μg/kg LPS (inflammation) for 24 h (n=8-11 animals per group). Violin plots (median and quartiles) depict microglial cell numbers. *P<0.05 (Two-way ANOVA). **I**, Histological confocal analysis of CD68 and Iba1 on tissue sections from neocortices of Rac1^fl/fl^ and Cx3cr1^CreER+^:Rac1^fl/fl^ mice after treatment with saline (CT) or intraperitoneal injection of 4 μg/kg LPS (inflammation) for 24 h (n=108 cells pooled across 4 animals per group). Violin plots (median and quartiles) depict CD68 fluorescence per Iba1^+^ cell. ****p<0.0001 (Two-way ANOVA). Scale bar: 20 μm.

To corroborate the role of Rac1 in modulating microglial sensing of external cues in vivo, we evaluated microglial activation in the brains of Cx3cr1^CreER+^:Rac1^fl/fl^ and Rac1^fl/fl^ littermates using LPS as a paradigm for brain inflammation. As expected, following brain inflammation (triggered by a single systemic injection of LPS), we observed, using immunofluorescence on tissue sections and flow cytometry, a robust microgliosis in Rac1^fl/fl^ brains (**Fig. 2G** and **H**); however, ablation of microglial Rac1 entirely prevented the inflammation-induced microgliosis in Cx3cr1^CreER+^:Rac1^fl/fl^ brains (**Fig. 2G** and **H**). We also evaluated changes in the expression of CD68 (a phagocytic marker highly expressed by activated microglia ^32^). We found that LPS-induced brain inflammation substantially increased the amounts of CD68 in Iba1^+^ cells in the brains of Rac1^fl/fl^ mice, but this effect was significantly attenuated in Iba1^+^ cells from Cx3cr1^CreER+^:Rac1^fl/fl^ brains (**Fig. 2I**). These results confirm the RNA-seq data showing that microglial environmental sensing requires Rac1 signaling.

### Rac1 regulates microglia-synapse crosstalk

The second core transcriptomic program — microglia-synapse crosstalk — also included two modules: intracellular protein trafficking, sorting, and export/secretion, and ECM/Growth factor signaling (**Fig. 1F** and **Suppl. Table 5**). The protein trafficking/export/secretion module included pathways related to Rho, Ras, and Rab GTPases, Rab-dependent protein trafficking, and cargo flux through microglial endomembranes. The ECM/Growth factor signaling module included adhesion/extracellular matrix (ECM), GDNF/NCAM-dependent pathway, EPH-ephrin signaling, and synaptic function (**Fig. 1F** and **Suppl. Table 5**). Microglia critically remodel the ECM and secrete growth factors that alter synaptic plasticity; thus, the RNA-seq results suggest that Rac1 supports a microglia-dependent growth factor signaling at synaptic interfaces.

Glial cell-derived neurotrophic factor (GDNF) interacts with the neuronal ECM (via NCAMs) ^33^ and ephrins ^34^ to modulate synapse formation and plasticity ^35-37^, and is sorted in and secreted from vesicles enriched in Rab GTPases ^38^. Thus, instructed by our RNA-seq results, we reasoned that Rac1 could modulate GDNF content in microglia. Accordingly, Rac1 and GDNF colocalized in brain microglia (**Suppl. Fig. 3A**) and microglia in culture (**Suppl. Fig. 3B**). Using flow cytometry, we observed decreased protein amounts of GDNF in microglia from Cx3cr1^CreER+^:Rac1^fl/fl^ mice compared to microglia from Rac1^fl/fl^ mice (**Suppl. Fig. 3C**). Moreover, whereas the knockdown of Rac1 decreased the GDNF content (corroborating the flow cytometry data in Cx3cr1^CreER+^:Rac1^fl/fl^ brains), switching on Rac1 activation (using chemogenetically-controlled Rac1 activity ^39^) increased the GDNF content in microglia in culture (**Suppl. Fig. 3D** and **E**, respectively).

The mammalian Rab family comprises more than 60 genes, and the proteins of the Rab27/3 branch regulate vesicular trafficking and exocytosis ^40^. Rab27a is present in microglia in the brain, while neurons highly express Rab3a ^41^. Using immunofluorescence, we observed that Rab27a colocalized with Rac1 in microglia in the brain (**Suppl. Fig. 3F**). In addition, microglia KD for Rac1 had significantly less Rab27a^+^ puncta than control microglia, indicating that Rac1 modulates the microglial Rab27a content (**Suppl. Fig. 3G**). Furthermore, we confirmed that GDNF colocalized with Rab27a in microglia in the brain (**Suppl. Fig. 3H**) and microglia in culture (**Suppl. Fig. 3I**). Like Rac1 KD, knocking down Rab27a decreased the GDNF content in microglia in culture (**Suppl. Fig. 3J**). Thus, these results further support the transcriptomic data related to protein sorting (**Fig. 1F**) and suggest that Rac1 controls Rab27a-mediated GDNF secretion by microglia.

### Microglial Rac1 is required for synaptic homeostasis, learning, and memory

To show that microglial Rac1 directly impacts microglia-to-synapse communication, we looked for global effects in synaptic structure and density. We found that the brains of Cx3cr1^CreER+^:Rac1^fl/fl^ mice had reduced spine density (**Fig. 3A**) and decreased excitatory (PDS-95^+^/vGlut1^+^) synapse number (**Fig. 3B**). The decrease in the numbers of spines and excitatory synaptic puncta caused by disrupting microglia-to-synapse communication should alter cognitive function, including learning and memory. Thus, we used the Morris Water Maze test to evaluate learning and memory. Indeed, Cx3cr1^CreER+^:Rac1^fl/fl^ mice had a significant delay finding the hidden platform in the learning phase (**Fig. 3C**) and spent significantly less time in the target quadrant in the probe trial (**Fig. 3D**). We concluded that Rac1-dependent disruption of homeostatic microglia-to-synapse communication leads to synaptic structural changes that correlate with deficits in learning and memory.

**Figure 3.**
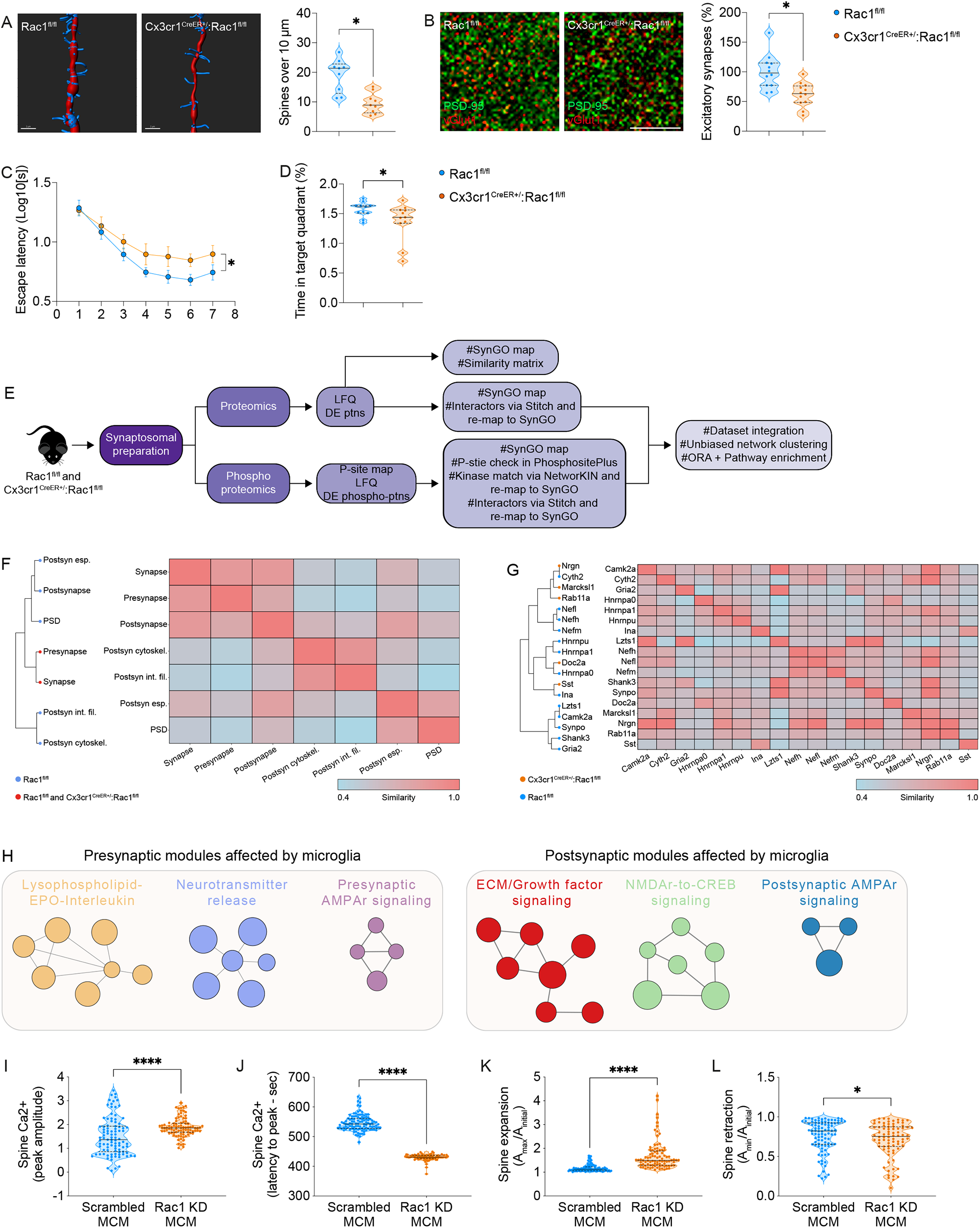
Microglial Rac1 is critical for synaptic homeostasis. **A and B**, Imaris rendering of dendritic spines and histological confocal colocalization of PSD-95 and vGlut1 on tissue sections from neocortices of Rac1^fl/fl^ and Cx3cr1^CreER+^:Rac1^fl/fl^ mice (n=9 dendrites stacked across 3 animals per genotype (A) or n=12 slices pooled across 4 animals per genotype (B)). Violin plots (median and quartiles) depict spine density or PSD-95/vGlut1 double-positive puncta. *P<0.05 (nested t-test (A); Mann-Whitney test (B)). Scale bars: A: 2 μm; B: 5 μm. **C and D**, Performance of Rac1^fl/fl^ and Cx3cr1^CreER+^:Rac1^fl/fl^ mice in the Morris Water Maze (n=11 mice per genotype). Histogram (mean and SEM) shows the learning curves, and violin plots (median and quartiles) depict the probe tests. *p<0.05 (two-way ANOVA (C); unpaired t-test (D)). **E**, Workflow of proteomic and phosphoproteomic profiling in brain synaptosomes and dataset integration. **F and G**, Hierarchical clustering and similarity matrix heatmap of differentially expressed synaptic protein products (H) or enriched cellular component GO terms (G) from Rac1^fl/fl^ and Cx3cr1^CreER+^:Rac1^fl/fl^ synaptosomes. Scales: Low (0) to high (1) similarly score colored from light blue to light coral. **H**, Pre and postsynaptic functional modules controlled by microglial Rac1 signaling revealed by proteomic and phosphoproteomic integration (n=4-5 mice per genotype). Nodes are scaled according to enrichment scores. **I-L**, Primary cortical neuronal cultures co-expressing GCamp6f and Lifeact-DsRed were recorded in saline for 10 min and then recorded upon stimulation with conditioned media from primary microglial cultures (MCM) carrying scrambled sequence or Rac1 shRNA. In all conditions, neurons were co-stimulated with 20 mM KCl. Violin plots (median and quartiles) show single-spine time-lapse fluorescence changes (n=88-112 spines pooled across 3 independent neuronal cultures). *p<0.05; ****p<0.0001 (Mann-Whitney test).

### Mapping the synaptic proteomic network controlled by microglial Rac1

To understand the modifications in the synaptic signaling elicited by microglia, we conducted quantitative high-throughput proteomics and phosphoproteomics in synaptosomal preparations from the brains of Cx3cr1^CreER+^:Rac1^fl/fl^ and Rac1^fl/fl^ mice (**Fig. 3E**). Label-free quantification (LFQ) identified 4641 unique proteins from which 126 had their abundances differentially expressed (DE), comparing the proteomes of Cx3cr1^CreER+^:Rac1^fl/fl^ and Rac1^fl/fl^ synaptosomes (**Suppl. Table 5**). Moreover, phosphoproteomics analysis identified 4389 phosphopeptides, from which 210 (corresponding to 167 unique proteins) were significantly altered between Cx3cr1^CreER+^:Rac1^fl/fl^ and Rac1^fl/fl^ synaptosomes (**Suppl. Table 6**).

To interrogate the synaptic protein network regulated by microglia, we developed functional similarity matrices (**Fig. 3F** and **G**), allowing direct global analysis of similarity alterations in the proteome structure of Cx3cr1^CreER+^:Rac1^fl/fl^ and Rac1^fl/fl^ synaptosomes. We found 2 enriched Cellular Component ontology terms for Cx3cr1^CreER+^:Rac1^fl/fl.^ and 7 for Rac1^fl/fl^ synaptosomes. GO:0045202 (synapse) and GO:0098793 (presynapse), common to Cx3cr1^CreER+^:Rac1^fl/fl^ and Rac1^fl/fl^ mice, showed a higher similarity and differentiated themselves for the other postsynaptic clusters (**Fig. 3F**). We also found 13 unique proteins within Rac1^fl/fl^ proteome and 5 within Cx3cr1^CreER+^:Rac1^fl/fl^ proteome and constructed a comparable similarity matrix for the DE synaptosomal proteins (**Fig. 3G**). These results indicate a substantial loss of functional modular polarization in the synaptic proteome of Cx3cr1^CreER+^:Rac1^fl/fl^ mice.

Then, we used an ontology-based pipeline (**Fig. 3E**) to map the synaptic signaling regulated by microglia. Network clustering segregated each synaptic compartment into three distinct functional modules (**Fig. 3H**). Whereas the presynaptic modules regulated by microglia included neurotransmitter release, presynaptic AMPA receptors, lysophospholipid/sphingosine 1-phosphate receptor, erythropoietin, and interleukin signaling, the postsynaptic modules included AMPA receptor activation and synaptic plasticity, NMDA-depended Camk2/CREB activation, and ECM/growth factor regulation of synaptic signaling. Various synaptic proteins altered in a microglia-dependent manner were common to both pre and postsynaptic compartments (**Suppl. Fig. 4** and **Suppl. Table 7**), including the major integrators Camk2a, Gria2, Efnb3, and Gephyrin (Gphn). Synaptic proteins regulated by microglia exclusive to the presynaptic network included the synaptic vesicle fusion regulator Doc2a, and the modulators of presynaptic vesicle clustering/cycling/priming Stxbp5, Sv2a, Syn1, Nsf, Rph3a, Bsn, and Pclo (**Suppl. Fig. 4** and **Suppl. Table 7**). Synaptic proteins regulated by microglia exclusive to the postsynaptic network included the critical postsynaptic scaffolds/organizers Shank3, Ctnnd2, PSD-93, Sptbn1, and Nrgn, and the critical synaptic activity integrators, DARPP-32, Rgs14, and Erk2 (**Suppl. Fig. 4** and **Suppl. Table 7**).

Altogether, these results raise the hypothesis that synaptic remodeling requires microglial Rac1. To test this hypothesis, we incubated primary cortical neurons with conditioned media obtained from control (scrambled MCM) or Rac1-deficient (Rac1 KD MCM) microglia and evaluated single-spine actin cytoskeleton and Ca^2+^ dynamics during activity-dependent spine remodeling (**Suppl. Fig. 5A**). Single-spine Ca^2+^ imaging (using the Ca^2+^ sensor GCaMP6f) revealed largely different kinetics of activity-dependent Ca^2+^ increase between scrambled MCM spines (**Suppl. Fig. 5B;** 1/slope = 595 au.s^-1^) and Rac1 KD MCM spines (**Suppl. Fig. 5B;** 1/slope = 2099 au.s^-1^). Such different time-dependent kinetics correlated with higher (**Fig. 3I** and **Suppl. Fig. 5C**) and faster (**Fig. 3J** and **Suppl. Fig. 5C**) activity-dependent Ca^2+^ responses in Rac1 KD MCM spines, indicating that loss of Rac1 signaling in microglia leads to aberrant spine Ca^2+^ overload during synaptic activity.

It is widely accepted that spine Ca^2+^ dynamics modulate spine structural changes. Thus, we investigated the remodeling of scrambled MCM spines and Rac1 KD MCM spines during synaptic activity. In both conditions, the overall dynamics of activity-dependent spine remodeling were similar (**Suppl. Fig 5D**), with an initial fast and transitory spine enlargement alongside actin polymerization followed by sustained spine contraction paralleled by actin depolymerization. However, the kinetics of activity-dependent remodeling varied substantially between scrambled MCM spines and Rac1 KD MCM spines (**Fig. 3K and L** and **Suppl. Fig 5E-N**). Specifically, Rac1 KD MCM spines grew bigger and faster during the enlargement phase (**Fig. 3K** and **Suppl. Fig. 5E**). Rac1 KD MCM spines also shrank more and faster during the contraction phase (**Fig. 3L** and **Suppl. Fig. 5F**).

Furthermore, in scrambled MCM spines, the activity-dependent spine enlargement and spine actin polymerization were desynchronized (**Suppl. Fig. 5G**), whereas transitory spine enlargement was associated positively with spine Ca^2+^ increase (**Suppl. Fig. 5H**). Thus, the activity-dependent transitory enlargement of scrambled MCM spines depended largely on Ca^2+^-associated membrane elasticity and not actin cytoskeleton elongation, which is in line with an actin polymerization-independent mechanism for transitory spine enlargement during synaptic activity in hippocampal slices ^42^. However, in Rac1 KD MCM spines, both activity-dependent actin polymerization and Ca^2+^ increase positively associated with abnormal spine enlargement (**Suppl. Fig. 5I and J**), indicating that the lack of microglial Rac1 relay caused aberrant spine enlargement through Ca^2+^ overload.

Moreover, the association of Ca^2+^ and actin dynamics with activity-dependent spine contraction also differed markedly between scrambled MCM spines and Rac1 KD MCM spines (**Suppl. Fig. 5K-N**). In scrambled MCM spines, actin depolymerization and Ca^2+^ increase positively associated with activity-dependent spine contraction (**Suppl. Fig. 5K and L**), showing that, in normal conditions, spine Ca^2+^ increase during synaptic activity promotes spine shrinkage by driving actin depolymerization. The more significant and faster spine contraction found in Rac1 KD MCM spines during synaptic activity associated positively with actin depolymerization (**Suppl. Fig. 5M**) but not with spine Ca^2+^ dynamics (**Suppl. Fig. 5N**), suggesting that the initial Ca^2+^ overload in Rac1 KD MCM spines also drives disproportionate actin depolymerization leading to excessive spine shrinkage during synaptic activity. These data indicate that microglial Rac1 modulates synaptic remodeling triggered by neuronal activity.

The decrease in spine density/synapse numbers found in Cx3cr1^CreER+^:Rac1^fl/fl^ brains could result from loss of a growth factor-associated mechanism modulating homeostatic microglia-to-synapse communication. Therefore, we evaluated Ca^2+^ dynamics in Rac1 KD MCM spines supplemented with exogenous GDNF (**Suppl. Fig. 6A**). Indeed, supplementing the Rac1 KD MCM with GDNF prevented the aberrant spine Ca^2+^ overload (**Suppl. Fig. 6B**), indicating that microglia-derived GDNF is required for homeostatic Ca^2+^ dynamics during neuronal activity. Moreover, we have not observed significantly reduced neuronal cell numbers (**Suppl. Fig. 6C**), changes in the pruning of excitatory synapses (**Suppl. Fig. 6D**), or neuroinflammation (**Suppl. Fig. 6E**) in Cx3cr1^CreER+^:Rac1^fl/fl^ brains, further supporting a role for a growth factor-associated mechanism for microglia-to-synapse communication in these mutants.

### Microglial Rac1 drives experience-dependent synaptic proteome plasticity

Because the lack of Rac1 signaling in microglia caused marked synaptic changes in the steady-state brain, we hypothesized that it could also impact the synaptic landscape driven by experience. Experience-dependent remodeling is the capacity of synapses to change and adapt to environmental inputs and learning ^43^ and relies on multistep phosphorylation of synaptic proteins ^44^. Thus, we used the environmental enrichment (EE) paradigm, which relates to improving sensory, motor, and cognitive stimulation in the housing conditions, to drive context-dependent synaptic remodeling. Then, we used quantitative phosphoproteomics in hippocampal synaptosomal preparations obtained from Rac1^fl/fl^ and Cx3cr1^CreER+^:Rac1^fl/fl^ mice housed under EE or control environment (CE) (**Fig. 4A**) to elucidate changes in the remodeling of the synaptic signaling modulated by microglial Rac1.

After unbiased high-confidence mapping, we found that the synaptic phosphoproteome driven by EE featured modifications in the abundance of 622 phosphopeptides (out of 3602) corresponding to 390 phosphoproteins (**Fig. 4B**). From the 622 phosphopeptides, 298 emerged specifically during EE, with upregulation of other 303 and downregulation of 21 phosphopeptides (**Fig. 4B**). To extract consensual information related explicitly to synaptic pathways, we used ORA using the 390 matched phosphoproteins revealed that EE significantly modified over 600 biological processes, 152 molecular functions, and 206 cellular components in the synapses (**Fig. 4B** and **Suppl. Table 9**). Pre and postsynapse organization, SNARE complex assembly, secretion of neurotransmitters, GABA and glutamate-dependent synaptic signaling, and postsynaptic cytosolic calcium levels were among the top signaling pathways modulated by EE in Rac1^fl/fl^ hippocampus (**Fig. 4C** and **Suppl. Table 9**). Unbiased PPI clustering identified five distinct phosphoprotein hubs comprising the EE-driven synaptic phosphoproteome (**Fig. 4D**).

We then used set theory to segregate the 390 synaptic phosphoproteins into two functional modules regulated by EE (**Fig. 4E** and **Suppl. Table 10**): one utterly independent of microglial Rac1 (138 phosphoproteins) and another strictly modulated by microglial Rac1 (149 phosphoproteins). Coalescing ORA, gene ontology, and unbiased PPI clustering allowed us to map the experience-dependent synaptic signaling modules exclusively related to microglial Rac1 (**Fig. 4F and G** and **Suppl. Table 10**).

**Figure 4.**
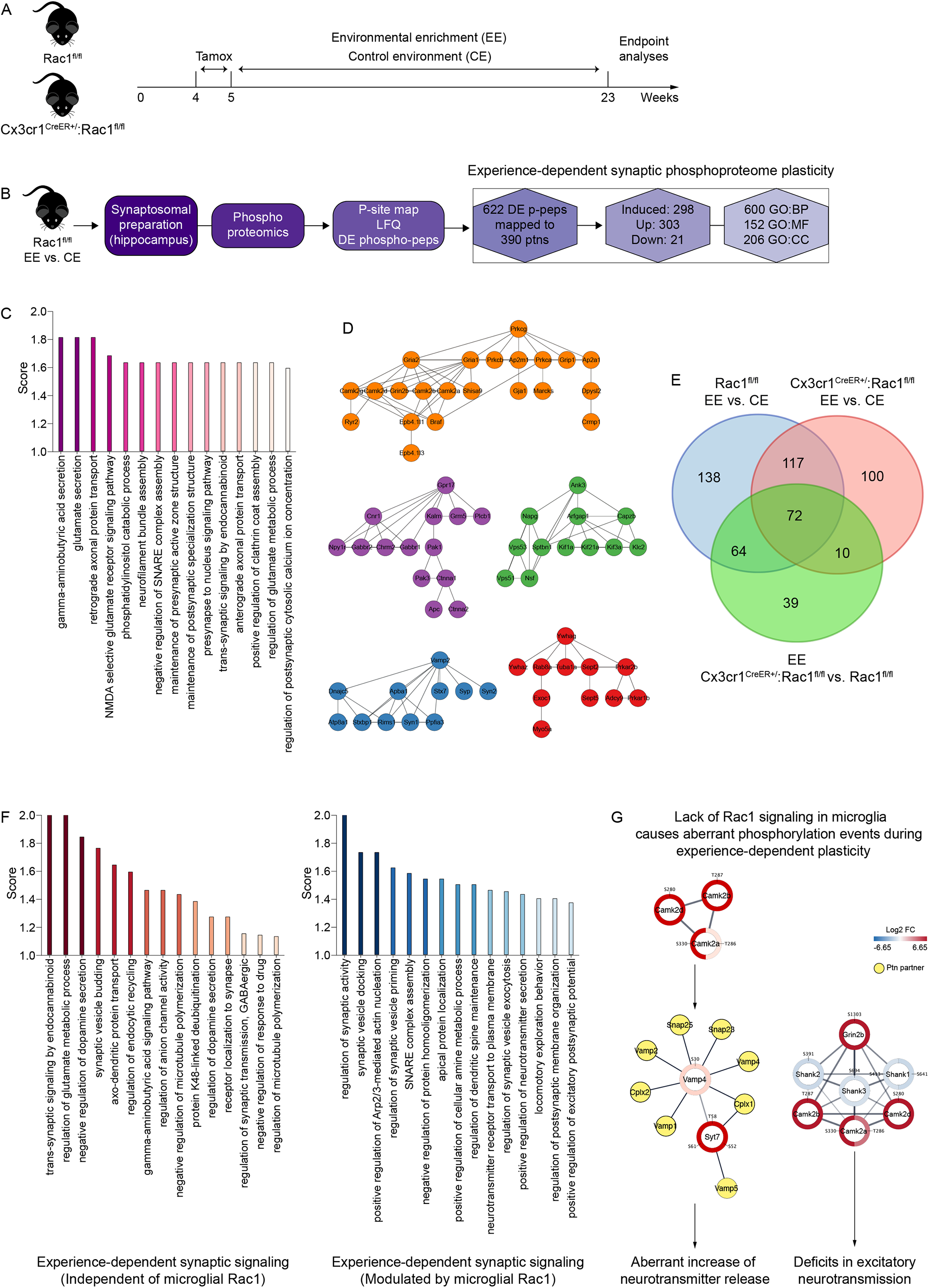
Microglial Rac1 modulates the phosphoproteome plasticity of the synapse. **A**, Housing conditions to elicit experience-dependent synaptic plasticity. **B**, Workflow of phosphoproteomic profiling in hippocampal synaptosomes after experience-dependent plasticity. **C**, Synaptic pathways modulated by experience (EE vs. CE) revealed by phosphoproteomic analyses (n=3-5 Rac1^fl/fl^ mice). The histogram displays the top enriched pathways. **D**, PPI hubs relative to EE vs. CE in Rac1^fl/fl^ mice. The top relevant phosphoproteins (based on C) altered by experience are displayed. **E**, Venn diagram allocating the differentially expressed phosphoproteins across housing conditions and genotypes. **F and G**, Segregation of synaptic pathways (F) and Camk2-associated phosphoprotein hubs (G) modulated concomitantly by experience and microglial Rac1 (n=3-5 mice per group). Histograms display the top enriched pathways (F). Hubs with Camk2-associated phosphoproteins are shown (G).

Specifically, we found that synaptic protein trafficking, endocannabinoid-dependent trans-synaptic signaling, glutamate/dopamine metabolism, GABA-dependent neurotransmission, and regulation of microtubule polymerization were among the top representative pathways controlled by EE independently of microglial Rac1 (**Fig. 4F red bars** and **Suppl. Table 10**). Importantly, global neurotransmitter release/recycle mechanisms, spine maintenance, postsynaptic architecture, excitatory neurotransmission, and regulation of actin nucleation were the most representative pathways modulated by EE through microglial Rac1 (**Fig. 4F blue bars** and **Suppl. Table 10**).

To provide further mechanistic insight into how microglial Rac1-dependent regulation of dynamic phosphorylation events was connected to synaptic function (for instance, neurotransmitter release and excitatory postsynaptic activity) during experience-dependent plasticity, we focused on the prominent synaptic integrator Camk2. After sequence analysis and literature curation, we identified a context-and microglial Rac1-dependent Camk2 circuitry operating simultaneously at both synaptic compartments (**Fig. 4G**). We detected synaptic hyperphosphorylation of canonical T286 in Camk2a and T287 in Camk2b, both known to induce kinase activation, in a microglial Rac1-regulated manner. Increased phosphorylation of S330, known to control binding to Camk2b, Camk2d, and Camk2g, was also modulated by microglial Rac1 (**Fig. 4G**). At the presynapse, microglial Rac1-mediated overactivation of Camk2a most likely disarrayed the fusion machinery for neurotransmitter release (composed of increased phosphorylated forms of Syt7 and Vamp4 and their synaptic partners), leading to an aberrant increase of neurotransmitter release (**Fig. 4G**). At the postsynapse, microglial Rac1-mediated overactivation of Camk2a, elicited by hyperactivity of GluN2B-containing NMDA receptors (hyperphosphorylated at S1303) and derangements of PSD scaffolds (illustrated by decreased phosphorylation of Shank1, 2, and 3), likely compromised the integration of postsynaptic potentials leading to deficits in excitatory neurotransmission (**Fig. 4G**). Here we concluded that microglial Rac1 modulates the remodeling of the synaptic phosphoproteome during experience-dependent plasticity.

### Microglial Rac1 instructs experience-dependent cognitive performance

Dynamic remodeling of protein phosphorylation networks is critical for cognitive function ^45^. So, we postulated that by modulating the remodeling of the synaptic signaling, microglial Rac1 should also impact experience-dependent cognitive performance. In such a way, we conducted different behavioral tests to evaluate the enhancement of the cognitive performance in Rac1^fl/fl^ and Cx3cr1^CreER+^:Rac1^fl/fl^ mice housed under EE or CE (**Fig. 5A**).

First, we assessed contextual memory in the step-down passive avoidance test ^46^. Using 3-Way ANOVA, we found, as expected, that the step-down latency was significantly higher in the test session than in the training session (**Fig. 5B**). Multiple comparisons within 3-Way ANOVA revealed an intergroup variation in memory recall (test session latency; **Fig. 5B**), suggesting that EE and CE might have affected memory retention differently across genotypes. Indeed, using a retention index transformation, we found that EE significantly increased retention in Rac1^fl/fl^ mice (**Fig. 5C**), whereas ablating microglial Rac1 abolished the EE-induced improvement in memory retention (**Fig. 5C**).

**Figure 5.**
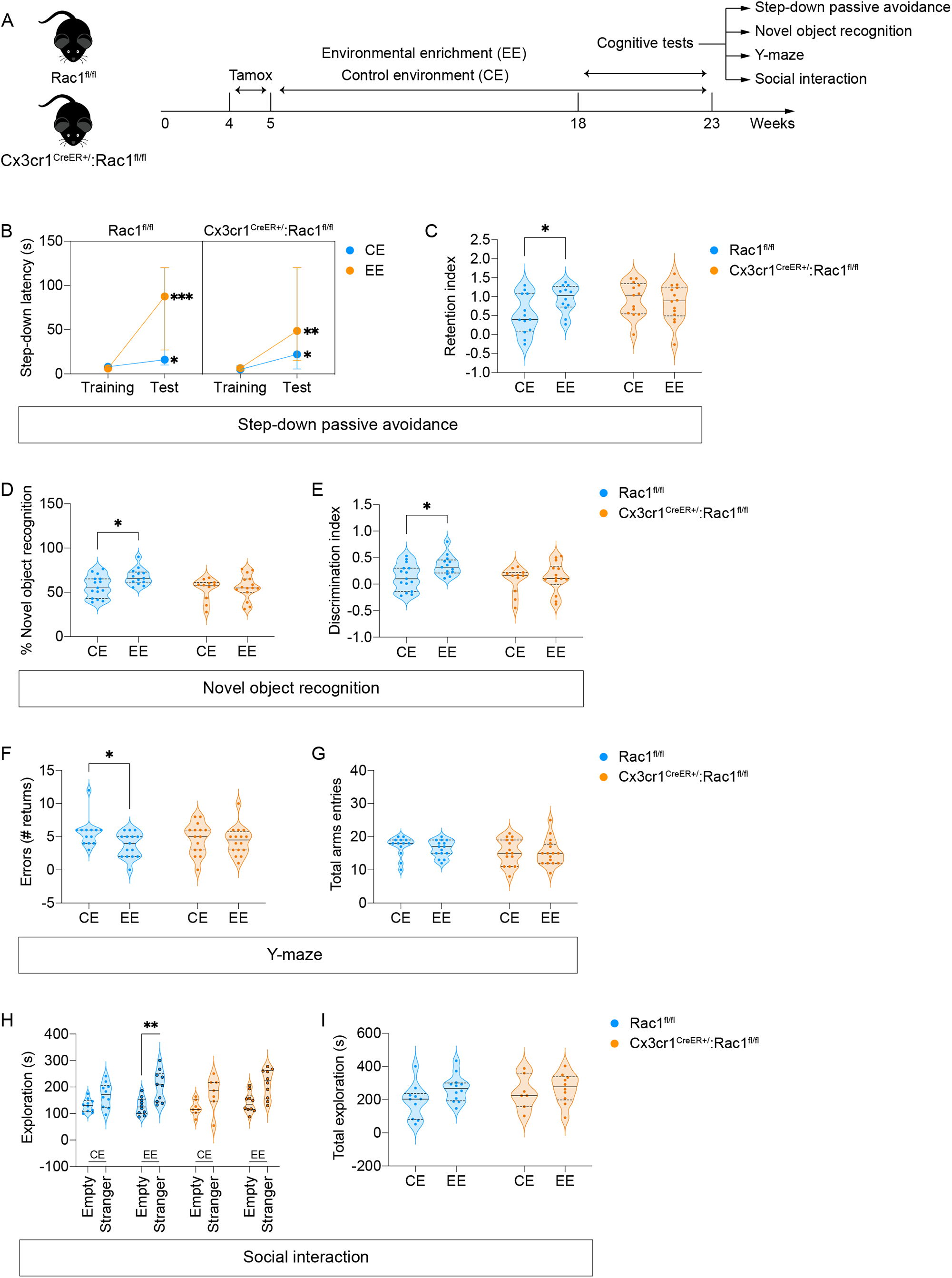
Experience-dependent cognitive performance requires microglial Rac1. **A**, Schematics for assessing cognitive performance driven by experience. **B-J**, Evaluation of Rac1^fl/fl^ and Cx3cr1^CreER+^:Rac1^fl/fl^ mice (n=13-16 mice (B-G) or 7-11 mice (H and I) per genotype) in the step-down passive avoidance (B and C), novel object recognition (D and E), Y-maze (F and G), and three-chamber social interaction tests (H and I). Histogram and violin plots display medians with quartiles. *p<0.05 (Three-way ANOVA (B); Two-way ANOVA (C-G); **p<0.01 and ***p<0.001 (Three-Way ANOVA).

Second, we used the novel object recognition (NOR) test to evaluate recognition memory ^47^ in Rac1^fl/fl^ and Cx3cr1^CreER+^:Rac1^fl/fl^ mice housed under EE or CE. We found that EE increased the degree of novel object exploration (**Fig. 5D**) and the discrimination of the novel object over the familiar object (**Fig. 5E**) in Rac1^fl/fl^ mice, indicating that EE improved recognition memory. On the other hand, the EE-induced improvement in the NOR performance was blocked entirely in Cx3cr1^CreER+^:Rac1^fl/fl^ mice (**Fig. 5D** and **E**).

Third, we used the Y-maze test to evaluate working memory ^48^ in Rac1^fl/fl^ and Cx3cr1^CreER+^:Rac1^fl/fl^ mice housed under EE or CE. We found that EE significantly decreased the number of arm entry errors, or incorrect arm returns, during the spontaneous maze alternation in Rac1^fl/fl^ mice (**Fig. 5F**), indicating that EE improves spatial working memory. However, the number of wrong arm returns was comparable between EE and CE in Cx3cr1^CreER+^:Rac1^fl/fl^ mice (**Fig. 5F**). As expected, no significant differences were found in total arms entries between groups among the genotypes (**Fig. 5G**). Here we concluded that the EE effect in enhancing spatial working memory requires microglial Rac1.

Lastly, we used the three-chamber social affiliation test ^49^ to evaluate social behavior in Rac1^fl/fl^ and Cx3cr1^CreER+^:Rac1^fl/fl^ mice housed under EE or CE. Whereas EE significantly increased Rac1^fl/fl^ mice’s preference for exploring the compartment with a stranger animal, EE failed to improve social proximity choices in Cx3cr1^CreER+^:Rac1^fl/fl^ mice (**Fig. 5H**). As expected, total exploration time was comparable among housing conditions and genotypes (**Fig. 5I**). Here we concluded that microglial Rac1 signaling modulates experience-dependent social interaction.

Overall, we concluded that microglial Rac1 instructs experience-dependent learning, memory, and sociability by modulating synaptic phosphoproteome plasticity.

## Discussion

Analysis of the RNA-seq profiling of Rac1-deficient microglia indicated that microglial sensing of external cues required Rac1 signaling. Combining functional in vitro and in vivo studies confirmed that Rac1 controls many signaling pathways critical for ligand-induced microglial modulation, including NF-kB modulation, ROS production, Ca2+ mobilization, and lipid sensing. Previous in vitro studies also show a Rac1 dependency for ROS generation, production of inflammatory mediators, and phagocytosis in microglia-like cells exposed to fibrilar amyloid ^53, 54^, further supporting the view that Rac1 is necessary for microglial environmental sensing.

Microglia are now recognized as critical for synaptic structure, function, and plasticity. Microglia-synapse crosstalk may occur through direct physical contact between microglia and synaptic elements ^55^, microglia-mediated ECM remodeling around synapses ^2^, or activation of synaptic receptors by microglia-secreted molecules (secretome) ^3, 4^. RNA-seq profiling of Rac1-deficient microglia revealed impaired intracellular protein trafficking and secretion. Additionally, loss of microglial Rac1 did not significantly affect the number of neurons, neuroinflammation, and engulfment capacity of synaptic elements. On this basis, we reasoned that the lack of Rac1 signaling could be modifying the microglial secretome profile to induce the synaptic and behavioral deficits observed. In agreement with this hypothesis, we identified in vivo the microglial Rab27a/GDNF (linked to the RNAseq module of protein trafficking and secretion) as a putative downstream target of Rac1. Moreover, the lack of GDNF signaling detrimentally affects synaptic function ^33, 35-37^ and spatial memory ^56^, which concur with the synaptic and behavioral alterations reported here for the Rac1 mutants.

GDNF can signal through NCAM to drive spine formation and postsynaptic assembly ^36^, whereas loss of NCAM decreases synapse numbers ^57^ and similarly disrupts spatial memory ^58^. A postsynaptic adhesion complex formed by NCAM, βI-spectrin, and NMDA receptors recruits and activates CamK2a to modulate synapse formation ^59^. Accordingly, the proteome and phosphoproteome profiling in brain synaptosomes of Rac1 mutants showed alterations in NCAM, spectrins, NMDA receptors, and CamK2a. On the other hand, we found no overt proteomic or phosphoproteomic evidence for a synaptic-related effect of brain-derived neurotrophic factor (BDNF) in microglial Rac1 mutants. The microglial BDNF phenotype relates more to an impairment in recognition memory and deficits in the rate of synapse formation/elimination, mainly in the motor cortex ^3^.

Molecular remodeling of synapses allows the brain to continuously change and adapt to novel environmental contexts and relies mainly on multistep modifications of synaptic proteins, including the modulation of their expression ^60^ and phosphorylation ^44^. Microglia-specific ablation of Rac1 combined with high-throughput proteomics allowed us to map and specify a microglia-dependent synaptic phosphoproteome signature driven by experience. Moreover, the substantial overlap of synaptic phosphoproteins (our datasets) and many mRNA species present at synaptic sites (e.g., CamK2a, PKC, PKA, NMDA receptor subunits, and actin cytoskeleton-related)^61^ validates our synaptic proteome profiling and is in line with a role for microglia in modulating the neuronal transcriptome ^2^. As such, our data further support the emergent notion of proteome plasticity, in which the proteome signature of an organism adapts to long-term changes to the external environment ^62^.

The fact that microglia shaped the proteome plasticity of the synapses logically suggested that microglia also impacted cognition. We provided evidence that microglia also controlled experience-dependent cognitive enhancement by remodeling the synaptic proteome. In future studies, mapping the experience-dependent proteome signature of microglia will provide a deeper understanding of how microglia shape the synaptic landscape to support cognitive performance.

Synaptic remodeling and plasticity ultimately govern the encoding and storage of new experiences in the brain, consequently dictating the behavioral adaptations to different environmental contexts. We triggered context-dependent synaptic remodeling using a paradigm of environmental enrichment (EE). It is well accepted that the molecular and cellular changes that EE produces in the brain correlate with enhanced cognitive function ^63^.

Indeed, we found that control mice housed under EE displayed notorious improvements in learning, memory, and sociability, confirming that EE improved cognitive performance. However, EE failed to produce cognitive enhancement in microglial Rac1 mutants. The vast differences found in the phosphoproteome signature in hippocampal synapses of microglial Rac1 mutants at least partially explain why EE failed to function as a cognitive enhancer in these mutants. Hence, the combination of synaptic proteomics profiling and behavioral tests indicated that microglial function is required to trigger synaptic proteome plasticity connected to cognitive performance.

Overall, we established Rac1 as a critical integrator of microglial homeostatic functions in the adult brain. Conditional cell-specific gene ablation, RNA-seq profiling, and functional in vitro and in vivo studies identified microglial Rac1 as a relay switch for microglial environmental sensing and microglia-synapse crosstalk. Combining microglial Rac1 ablation with synaptosome proteome profiling allowed us to map for the first time a microglial Rac1-dependent synaptic proteomic signature driven by experience. Furthermore, integrating behavioral analyses in mice lacking Rac1 in adult microglia revealed a novel function for microglia as a modulator of experience-dependent cognitive enhancement. Finally, mapping the synaptic changes elicited by disrupting microglial homeostatic functions might have broad implications for devising strategies to enhance cognitive performance or compensate for the cognitive decline during aging or brain diseases.

### Experimental Procedures

#### Animals

All mice experiments were reviewed by i3S animal ethical committee and were approved by Direção-Geral de Alimentação e Veterinária (DGAV). Animals were maintained with an inverted 12h/12h light-dark cycle and were allowed free access to food and water. Mice were housed under standardized cages and specific pathogen-free conditions. Experiments were carried out following the 3Rs ethics policy.

##### Conditional Rac1-deficient mice

Cx3cr1^CreER-EYFP^ mice were purchased from Jackson Laboratories and used as before ^4^. In such mice, the *Cx3cr1* promoter drives high expression of the CreER cassette in microglia ^3^. Mice homozygous for the *Rac1* floxed allele ^16^ were backcrossed at least ten generations and were kept at the I3S animal facility. PCR determined all genotypes on genomic DNA. Rac1 floxed mice were crossed with Cx3cr1^CreER-EYFP^ mice. Progeny of interest were: Control (Rac1^fl/fl^ or Cx3cr1^CreER+^) and mutants (Rac1^fl/fl^:Cx3cr1^CreER+^). Mice (4/5-weeks-old) were given tamoxifen (5 mg *per* animal by oral gavage; 1 mg daily for five days) and then analyzed 6-8- or 18-weeks post-tamoxifen. Experiments were performed in female and male mice (unless specified otherwise), all kept on a C57BL/6 background.

#### Antibodies

The following antibodies were used in this study: Rac1 antibody [0.T.127] (Abcam Cat# ab33186, RRID:AB_777598; 1:250), Rab27a mouse monoclonal (M02), clone 1G7 (Abnova Cat# H00005873-M02, RRID:AB_519010; 1:200), Rab27a rabbit polyclonal (Sigma-Aldrich Cat# HPA001333, RRID:AB_1079730; 1:200), GDNF (Abcam Cat# ab18956, RRID:AB_2111379; 1:100-1:200), GFP (Abcam Cat# ab6673, RRID:AB_305643; 1:200), Iba1 (Wako Cat# 019-19741, RRID:AB_839504; 1:500), CD11b clone M1/70.15 (BD Biosciences Cat# 560456, RRID:AB_1645267; 1:100), PSD95 clone 6G6-1C9 (Thermo Fisher Scientific Cat# MA1-045, RRID:AB_325399; 1:600), vGlut1 (Synaptic Systems Cat# 135 303, RRID:AB_887875; 1:1,000), CD68 clone FA-11 (Bio-Rad Cat# MCA1957T, RRID:AB_2074849; 1:400), Fc Receptor Blocking Solution (BioLegend Cat# 156603, RRID:AB_2783137; 1:50), NeuN (Millipore Cat# MAB377, RRID:AB_2298772; 1:400), APC anti-mouse/human CD11b antibody (BioLegend Cat# 101212, RRID:AB_312795; 0.25 μg to 10^6^ cells), Alexa Fluor® 647 anti-mouse/human CD11b antibody (BioLegend Cat# 101218, RRID: AB_389327; 0.25 μg to 10^6^ cells), PE anti-mouse CD45 antibody (BioLegend Cat# 103106, RRID:AB_312971; 0.25 μg to 10^6^ cells).

#### Primers

The primers used in this study were purchased from Thermo Fisher Scientific; their sequences are provided in **Suppl. Table 11**.

#### Plasmids

pFRET-HSP33 cys (RRID:Addgene_16076), IκBα-miRFP703 (RRID:Addgene_80005), Daglas-pm1 (PMID: 16990811), pcDNA-D1ER (RRID:Addgene_36325), CMV-R-GECO1 (RRID:Addgene_32444), CKAR (RRID:Addgene_14860), pLKO-empty (PMID: 28960571), pLKO-Rac1 shRNA (PMID: 28960571), pLKO-Rab27a shRNA (Sigma TRCN0000279985), scrambled-mCherry shRNA control (PMID: 28960571), mCherry-Rac1 shRNA (PMID: 28960571), pRK5-myc-Rac1-wt (RRID:Addgene_12985), pRK5-myc-Rac1-T17N (RRID:Addgene_12984), YF-Rac1(CA) (RRID:Addgene_20150), Lyn11-targeted FRB (LDR) (RRID:Addgene_20147), YFP-FKBP (YF) (RRID:Addgene_20175) pLKO.1 (Sigma), pLKO-Rab27a shRNA (Sigma), mCherry-Lifeact-7 (RRID:Addgene_54491), pGP-CMV-GCaMP6f (RRID:Addgene_40755).

#### Environmental enrichment protocol

Mice with 4-5 weeks of age were weighted and placed in Type III cages three days with two different housing conditions: control environment (CE) and enriched environment (EE). Littermates were randomly assigned to one of the housing conditions, with Rac1^fl/fl^ and Cx3cr1^CreER+^:Rac1^fl/fl^ mice in every cage. CE only contained bedding material (corn cob, grade 12) and tissue paper. EE had bedding material, tissue paper, a small cardboard tube, four small aspen bricks, a wood ladder, a vertical plastic running wheel with a steep metal grid, a dome home, a plastic Lego-like block, and sunflower seeds. Mice were weighted immediately before tamoxifen administration and then weighted once per week during home-cage change for the first three weeks after tamoxifen administration. Mice were kept in the same housing conditions until the end of the protocol (18 weeks) with access to food and water *ad libitum*. Home-cages in both CE and EE were changed weekly. Enrichment material was checked every week: new tissue paper and sunflower seeds were added weekly, while cardboard tubes, dome homes, and aspen bricks were replaced when destroyed or highly degraded. The remaining enrichment material was constantly transferred to the new cage during weekly home-cage changes. The spatial organization of the enrichment material was maintained throughout the experiment. Manipulation of mice during the behavioral test period was reduced to the minimum necessary.

#### Behavioral tests

Procedures were conducted in the dark phase of the light/dark cycle and performed blind to genotypes. The order and interval between tests were used as acclimation for the next test and performed in the following order: (1) three-chamber sociability; (2) novel object recognition; (3) Y-maze; (4) step-down passive avoidance. The Morris water maze test was conducted in separate and independent animal cohorts. The experimenter recording the behavioral parameters during test sections was blind to genotype.

##### Three-chamber sociability

Mice were first habituated to the empty apparatus, a three-chamber box consisting of three interconnected lined compartments, for 10 minutes. After the habituation phase, mice were tested in the sociability task. In this phase, subject mice socialize with a conspecific mouse in a cage or explore an empty cage in opposite external compartments. All phases during 10 minutes and the placement of stranger mice on the left or right side of the chamber are systematically altered between trials. The time spent in each compartment (four paws have entered the chamber) and the exploration time (direct contact or stretching in an area of 3 – 5 cm around the mouse or empty cage) were measured. All parameters were evaluated by Boris software ^64^. Because female control mice did not significantly enhance sociability parameters, only male mice (both genotypes) were used in this test. Statistical analyses were conducted using three-way ANOVA (exploration times in empty and stranger compartments were fixed as row factors) or randomized-block two-way ANOVA (mice were considered the experimental units; statistical significance was defined as P < 0.01) with sphericity assumption and the Turkey correction for multiple comparisons.

##### Object recognition (NOR)

The NOR test used experimental procedures similar to the three-chamber test. The test consists of three phases. During the habituation phase, mice can explore the apparatus for 10 min (used to perform habituation in three-chamber test, 24 h before). The next day, during the object familiarization/acquisition phase, two identical objects were placed at the center of each outer chamber. Mice were allowed to explore the objects for 10 min freely. Then mice were returned to their home cage, and after 4 h (inter-trial interval, ITI), the retention phase was performed. In this phase, one of the familiar objects is changed by a novel object, and animals are allowed to explore these objects for 3 min. Exploration was defined as follows: the mouse touched the object with its nose, or the mouse’s nose was directed toward the object at a distance shorter than 2 cm. The discrimination index (DI) was calculated as before ^4^ and used as an index of memory function, DI = (time exploring the novel object) / (total time spent exploring both objects). All parameters were evaluated by Boris software ^64^. Statistical analyses were conducted using randomized-block two-way ANOVA (mice were considered the experimental units; statistical significance was defined as P < 0.05) with sphericity assumption and the Turkey correction for multiple comparisons.

##### Y-maze

Wrong arm returns (# errors), a measure of spatial working memory, was assessed by allowing mice to explore all three arms of the maze, motivated by an innate interest of rodents to explore previously unvisited areas. The mice are positioned in the center of the apparatus and allowed to explore freely for 8 minutes. During the test, visual clues were placed on the walls. Entries in the same arm during the spontaneous alternation phase were considered wrong arm returns and used as the number of working memory errors. Statistical analyses were conducted using randomized-block two-way ANOVA (mice were considered the experimental units; statistical significance was defined as P < 0.05) with sphericity assumption and the Turkey correction for multiple comparisons.

##### Step-down passive avoidance

The step-down test was used to assess the long-term memory and consisted of two phases. First, in the training phase, each mouse was placed in the center of an elevated platform (15 mm above the grid floor), and the step-down from the platform with four paws was immediately followed by a foot-shock (0.5 mA) for 2 s. The latency to step-down was measured (maximum of 120s; minimum 10 s). Afterward, the animals were presented to the retention phase 24 h after training. This phase was conducted in the absence of shock, and the step-down latency from the platform was recorded (up to 120 s) and evaluated as indicators of memory retention. The retention index was calculated using the step-down latency from the test session divided by the latency from the training session. Statistical analyses were conducted using three-way ANOVA (step-down latencies in training and retention were fixed as row factors) or randomized-block two-way ANOVA with sphericity assumption. The Turkey correction for multiple comparisons was conducted in both analyses. Moreover, mice were considered the experimental units, and statistical significance was defined as P < 0.05.

##### Morris water maze (MWM)

The MWM was used to evaluate spatial memory. The apparatus consisted of a circular pool (110 cm diameter) filled with water (21±1ºC) made opaque gray. Visual cues were positioned equidistant in the walls. Cued learning was performed during the first two days. A non-visible escape platform (7 × 8 cm) was submerged 1 cm below the water surface in the quadrant center. In this phase, mice were trained to find the hidden platform with a visual clue. Animals were subjected to four consecutive swimming trials with different start and goal positions. In the acquisition phase (4 trials/day for five days), mice were given up to 60 seconds per trial to find the hidden platform and were required to remain seated on the platform for 10 s. The platform location is the same during the acquisition phase (center of target quadrant). Escape latency (i.e., time to locate the platform) was measured during the cue learning and acquisition phase. During the probe test (day 8), the platform was removed from the pool, and each mouse was given up to 30 s to search the platform’s position. All parameters were automatically evaluated by SMART v3.0 software (Panlab, Barcelona, Spain). Statistical analyses to assess learning were conducted using repeated measures two-way ANOVA (mice were considered the experimental units; statistical significance was defined as P < 0.05) with the Greenhouse-Geisser correction (to combat sphericity violation). In the probe test, statistical analyses were conducted using unpaired t-test (mice were considered the experimental units; statistical significance was defined as P < 0.05).

#### Adeno-associated virus (AAV) injections and quantification of dendritic spines

Mice were anesthetized by isoflurane inhalation and positioned on a stereotactic frame. Mice eyes were lubricated to prevent cornea dryness. The animal was maintained with a Stoelting™ Rodent Warmer device with a rectal probe to keep its temperature at 37ºC. Deep anesthesia was confirmed by the absence of a response to a toe pinch. We used the following coordinates to target the hippocampus: anteroposterior (AP) -2.3, mediolateral (ML) ±1.3 mm, dorsoventral (DV) -2.2 mm, using the bregma as reference. Only the right hippocampus of each animal was injected with pAAV-hSyn-mCherry (Addgene viral prep # 114472-AAV5). The viral titer was 7×10^1 2^ vg/mL. The red fluorescent protein (mCherry) was driven by the synapsin-1 promoter.

The surgical procedure started with a skin incision in the mouse’s head to expose the bregma. After the correct X: Y localization, a drill was used to puncture the skull. The needle was slowly introduced into the animal’s brain until it reached the right z coordinate. We waited 2 minutes for tissue accommodation before the AAV administration. To optimize neuronal labeling to subjacent cortical areas, we injected 0.5 μl of undiluted viral preparation (corresponding to 7×10^1 2^ viral genomes (vg)) into the right hippocampus at a rate of 0.1 Lμl/min. The needle was removed 2 min after administration, the skin was sutured, and isoflurane flow stopped. After waking up, mice were put in a new cage for the first 24 hours, with wet food in the cage bedding, paracetamol in the water bottle, and dietary supplements (anima strath and Duphalyte®) to avoid weight loss and malnutrition.

Dendritic spines were reconstructed using Imaris. Briefly, dendrites were traced with the filaments tool using the AutoPath algorithm. The dendrite diameter was rebuilt using the shortest distance from the distance map algorithm with the local contrast threshold. Spines were detected with seed points set to 0.3 μm and the maximum spine length at 3 μm. The spine diameter was rebuilt using the shortest distance from the distance map algorithm with the local contrast threshold. Statistical analyses were conducted using nested t-test (to combat pseudoreplication) with spine density per dendrite stacked within each mouse (statistical significance was defined as P < 0.05).

#### Tissue preparation and immunofluorescence

After animal perfusion with ice-cold PBS (15 ml), brains were fixed by immersion in 4% PFA in PBS, pH 7.2 overnight. After that, brains were washed with PBS and then cryoprotected using a sucrose gradient in a row (15 and 30%). After 24 h, brains were mounted in an OCT embedding medium, frozen, and cryosectioned in the CM3050S Cryostat (Leica Biosystems). Coronal sections from brains (30 μm thickness) were collected non-sequentially on Superfrost ultra plus slides. Tissue sections from controls and experimental mice encompassing identical stereological regions were collected on the same glass slide and stored at -20°C. Frozen sections were defrosted for at least 1 hour and hydrated with PBS for 15 min. Sections were permeabilized with 0.25% Triton X-100 for 15 min, washed with PBS for 10 min and blocked (5% BSA, 5% FBS, 0.1% Triton X-100) for 1 hour. Primary antibodies were incubated in a blocking solution in a humidified chamber overnight at 4°C. Secondary antibodies were incubated for 2 hours in a blocking solution. After the secondary antibody, sections were washed three times for 10 min with PBS. Slides were coverslipped using glycergel or Immumount.

#### Confocal imaging and morphometric analysis

Images from tissue sections of the neocortex and the dorsal hippocampus were acquired in 8-bit sequential mode using standard TCS mode at 400 Hz. The pinhole was kept at one airy in the Leica TCS SP8 confocal microscope. Images were illuminated with different laser combinations and resolved at 512 × 512 or 1024 × 1024 pixels format using HyD detectors, and entire Z-series were acquired from tissue sections. Equivalent stereological regions were obtained for all tissue sections within a given slide.

To quantify microglia (Iba1+ cells) or neurons (NeuN+ cells), the number of cells was manually scored, as before ^4, 65^, in stereological identical regions of the neocortex of stained sections (3 images per section; 3 sections per mice for each experimental group). Statistical analyses comparing two genotypes were conducted using the Mann-Whitney test (mice were considered the experimental units; statistical significance was defined as P < 0.05). Statistical analyses comparing the effect of LPS between genotypes were conducted using two-way ANOVA (tissue sections were considered the experimental units; statistical significance was defined as P < 0.05) with the Šidák correction for multiple comparisons.

To quantify synapses, images from stereological identical neocortical regions from each experimental group (3 images per section; 3 sections per animal for each experimental group) were acquired using a Leica HC PL APO CS 40x /1.10 CORR water objective at 1024 × 1024 pixels resolution with 8-bit bidirectional non-sequential scanner mode at 400 Hz and pinhole at one airy in the Leica TCS SP5 II confocal microscope. Z-stacks were converted to maximum projection images using the LAS AF routine. Z-projections were background-subtracted using the rolling ball background subtraction built-in algorithm in FIJI, and then images were upsampled using a bicubic interpolation routine. The double-positive PSD-95/vGlut1 puncta per μm^2^ were manually scored for each image. Statistical analyses comparing two genotypes were conducted using the Mann-Whitney test (tissue sections (three per mice) were considered the experimental units; statistical significance was defined as P < 0.05).

To assess microglial morphology, the ramification of neocortical Iba1^+^ cells was evaluated as described before ^16^. Branches were traced using the NeuronJ plugin, and data for each cell were converted into SWC format using Bonfire. Using Bonfire, individual segments were connected to NeuronStudio software and audited for any tracing errors. Sholl analysis was performed by drawing concentric circles around the cell body at defined radius increments. Branching data were extracted using a custom MATLAB routine. Representative microglial morphology images were generated using 3D rendering in Imaris as before ^65^. Statistical analyses were conducted using randomized-block two-way ANOVA (single microglia were considered the experimental units; statistical significance was defined as P < 0.05) with sphericity assumption and the Šidák correction for multiple comparisons.

Colocalization analyses were carried out in Imaris or FIJI using the Coloc2 plugin (https://imagej.net/Coloc_2). Statistical analyses comparing two genotypes were conducted using the Mann-Whitney test (mice or single microglia were considered the experimental units; statistical significance was defined as P < 0.05). Statistical analyses comparing four groups were conducted using randomized-block two-way ANOVA (single microglia were considered the experimental units; statistical significance was defined as P < 0.05) with sphericity assumption and the Šidák correction for multiple comparisons.

#### Flow cytometry and cell sorting

The following markers were used to characterize microglia in the samples: CD45 CD11b and Ly6C. Microglia were collected from the brains of control and mutant mice using density gradient separation as before ^4, 65^. Single-cell suspensions (5 × 10^5^ cells) were incubated with different mixes of FACS antibodies for 30 min at 4°C in the dark. Compensation settings were determined using spleen from both control and mutant. Cell suspensions were evaluated on a FACS Canto II analyzer (BD Immunocytometry Systems). Cell suspensions were seeded in a U bottom 96-well plate. For GDNF staining, cells were treated with Brefeldin A (10 μg/mL) for 3 h at 37ºC in RPMI supplemented with 1% Pen/Strep and 10% FBS. Cells were incubated with CD45-PE and CD11b-Alexa Fluor 647. After antibody washing, for intracellular staining, cells were fixed in 2% PFA for 30 min, washed in PBS, and permeabilized with permeabilization buffer (Life Technologies 00-8333-56). Intracellular antibody staining mix was prepared in permeabilization buffer and incubated with the cells for 1 h at 4ºC in the dark. After washing in permeabilization buffer, cells were incubated with Alexa Fluor 488 secondary antibody for 30 min at 25ºC in the dark. After that, cells were washed twice in permeabilization buffer, resuspended in FACS staining buffer (2% BSA, 0.1% sodium azide), and analyzed in FACS Canto II. Antibody controls for Rac1 and GDNF staining (FMO) were prepared in mixes with CD45-PE and CD11b-Alexa Fluor 647 plus secondary antibodies alone. Cell sorting was performed on a FACS ARIA cell sorter as before ^4^. Data were analyzed by FlowJo X10 software (TreeStar).

Statistical analyses comparing two genotypes were conducted using the Mann-Whitney test (mice were considered the experimental units; statistical significance was defined as P < 0.05). Statistical analyses comparing three different genotypes were conducted using the Kruskal-Wallis test (mice were considered the experimental units; statistical significance was defined as P < 0.05) with the two-stage linear step-up procedure of Benjamini, Krieger, and Yekutieli correction for multiple comparisons. Statistical analyses comparing the effect of LPS between genotypes were conducted using two-way ANOVA (mice were considered the experimental units; statistical significance was defined as P < 0.05) with the Šidák correction for multiple comparisons.

#### MACS isolation of microglia

Mice (males) were perfused with ice-cold PBS, and their brains were removed. The right hemisphere was mechanically dissociated in ice-cold Dounce buffer (15mM HEPES; 0,5% Glucose; and DNAse) by six strokes in a tissue potter. Homogenate was pelleted by centrifugation, resuspended in MACS buffer (0.5% BSA; 2 mM EDTA in PBS) followed by incubation with 80 μL myelin removal microbeads. Homogenate was negatively selected using LS columns, pelleted, washed twice, resuspended in MACS buffer, and incubated 10 μL CD11b microbeads. CD11b^+^ fraction was selected using LS columns according to the manufacture’s instructions. Eluted CD11b-enriched fraction was centrifuged (9300 g; 1 min; 4ºC) and reserved for RNA isolation. RNA was isolated using the RNeasy Plus Micro Kit. RNA integrity was analyzed using the Bioanalyzer 2100 RNA Pico chips (Agilent Technologies, CA, USA).

#### Library preparation, Sequencing, and Bioinformatics

Ion Torrent sequencing libraries were prepared according to the AmpliSeq Library prep kit protocol as we did before ^66^. Briefly, 1 ng of highly intact total RNA was reverse transcribed. The resulting cDNA was amplified for 16 cycles by adding PCR Master Mix and the AmpliSeq mouse transcriptome gene expression primer pool. Amplicons were digested with the proprietary FuPa enzyme, and then barcoded adapters were ligated onto the target amplicons. The library amplicons were bound to magnetic beads, and residual reaction components were washed off. Libraries were amplified, re-purified, and individually quantified using Agilent TapeStation High Sensitivity tape. Individual libraries were diluted to a 50 pM concentration and pooled equally. Emulsion PCR, templating, and 550 chip loading were performed with an Ion Chef Instrument (Thermo Scientific MA, USA). Sequencing was performed on an Ion S5XL™ sequencer (Thermo Scientific MA, USA) as we did before ^66^.

Data from the S5 XL run processed using the Ion Torrent platform-specific pipeline software Torrent Suite v5.12 to generate sequence reads, trim adapter sequences, filter and remove poor signal reads, and split the reads according to the barcode. FASTQ and BAM files were generated using the Torrent Suite plugin FileExporter v5.12. Automated data analysis was done with Torrent Suite™ Software using the Ion AmpliSeq™ RNA plugin v.5.12 and target region AmpliSeq_Mouse_Transcriptome_V1_Designed as we did before ^66^.

Raw data was loaded into Transcriptome Analysis Console (4.0 Thermo Fisher Scientific, MA, EUA) and first filtered based on ANOVA eBayes using the Limma package and displayed as fold change. Significant changes had a p-value < 0.05 and FDR < 0.2. Functional enrichment analyses were performed using Gene Set Enrichment Analysis (GSEA) with WEB-based Gene Set Analysis Toolkit (WebGestalt) ^67^ and STRING ranked enrichment (SRE) ^68^. The whole transcriptome gene list was ranked (linear fold change*-log10 FDR) and submitted to WebGestalt (http://www.webgestalt.org) or STRING (https://string-db.org/cgi/input?sessionId=bFzQ6nknJDLN&input_page_show_search=on).

Pathway enrichment was performed using the REACTOME database, with default settings. Enrichment scores for gene sets were calculated using an FDR cutoff of 0.05. Enriched pathways were manually recategorized to core transcriptomic modules and are displayed as a network (constructed using Cytoscape). Contingency analyses covering different microglial phenotypes were performed using Fisher’s exact test and the Baptista-Pike method to calculate the odds ratio.

#### Gene expression by qRT-PCR

RNA was extracted from flow-cytometry sorted microglia using the Direct-zol™ RNA MiniPrep kit according to the manufacturer’s instructions. cDNA synthesis was performed using 300 ng of total RNA (DNase I treated) with SuperScript^®^ III First-Strand Synthesis SuperMix. qRT-PCR was carried out using iQ™ SYBR^®^ Green Supermix on an iQ™5 multicolor real-time PCR detection system (Bio-Rad). The expression of PCR transcripts was calculated using the 2^-deltaCt^ with *Yhwaz* serving as the internal control gene. Statistical analyses were performed on raw 2^-deltaCt^ values to detect differentially expressed transcripts between sampled groups. Statistical analyses were conducted using Mann-Whitney (mice were considered the experimental units; statistical significance was defined as P < 0.05).

#### Synaptosomal preparations

Synaptosomes were acutely prepared as before ^65^. One hundred micrograms of synaptosomal proteins from each sample were processed for proteomic analysis following the solid-phase-enhanced sample-preparation (SP3) protocol. Enzymatic digestion was performed with trypsin/LysC (2 micrograms) overnight at 37°C at 1000 rpm. The resulting peptide concentration was measured by fluorescence. Enrichment for phosphorylated peptides was performed using Titanium dioxide beads (TiO2; ThermoFisher Scientific) as described in the manufacturer’s protocol.

#### High-throughput proteomics data acquisition and quantification

Protein identification and quantitation were performed by nanoLC-MS/MS using an Ultimate 3000 liquid chromatography system coupled to a Q-Exactive Hybrid Quadrupole-Orbitrap mass spectrometer (Thermo Scientific, Bremen, Germany). Five hundred nanograms of peptides of each sample were loaded onto a trapping cartridge (Acclaim PepMap C18 100 Å, 5 mm × 300 μm i.d., 160454, Thermo Scientific, Bremen, Germany) in a mobile phase of 2% ACN, 0.1% FA at 10 μL/min. After 3 min loading, the trap column was switched in-line to a 50 cm × 75 μm inner diameter EASY-Spray column (ES803, PepMap RSLC, C18, 2 μm, Thermo Scientific, Bremen, Germany) at 300 nL/min. Separation was achieved by mixing A: 0.1% FA and B: 80% ACN, 0.1% FA with the following gradient: 5 min (2.5% B to 10% B), 120 min (10% B to 30% B), 20 min (30% B to 50% B), 5 min (50% B to 99% B), and 10 min (hold 99% B). Subsequently, the column was equilibrated with 2.5% B for 17 min. Data acquisition was controlled by Xcalibur 4.0 and Tune 2.9 software (Thermo Scientific, Bremen, Germany).

The mass spectrometer was operated in the data-dependent (dd) positive acquisition mode alternating between a full scan (*m*/*z* 380-1580) and subsequent HCD MS/MS of the 10 most intense peaks from a full scan (normalized collision energy of 27%). The ESI spray voltage was 1.9 kV. The global settings were as follows: use lock masses best (*m*/*z* 445.12003), lock mass injection Full MS and chrom. peak width (FWHM) of 15 s. The full scan settings were as follows: 70 k resolution (*m*/*z* 200), AGC target 3 × 10^6^, maximum injection time 120 ms; dd settings: minimum AGC target 8 × 10^3^, intensity threshold 7.3 × 10^4^, charge exclusion: unassigned, 1, 8, >8, peptide match preferred, exclude isotopes on, and dynamic exclusion 45 s. The MS2 settings were as follows: microscans 1, resolution 35 k (*m*/*z* 200), AGC target 2 × 10^5^, maximum injection time 110 ms, isolation window 2.0 *m*/*z*, isolation offset 0.0 *m*/*z*, dynamic first mass, and spectrum data type profile.

The raw data were processed using the Proteome Discoverer 2.5.0.400 software (Thermo Scientific, Bremen, Germany). Protein identification analysis was performed with the data available in the UniProt protein sequence database for the *Mus Musculus* Proteome (2020_02 version, 55,398 entries) and a common contaminant database from MaxQuant (version 1.6.2.6, Max Planck Institute of Biochemistry, Munich, Germany). Sequest HT tandem mass spectrometry peptide database search program was used as the protein search algorithm. The search node considered an ion mass tolerance of 10 ppm for precursor ions and 0.02 Da for fragment ions. The maximum allowed missing cleavage sites was set as 2. For the phosphoproteomics, the IMP-ptmRS node, with the PhosphoRS mode (set to false), was used to localize phosphorylation sites. The Inferys rescoring node was considered, and the processing node Percolator was enabled with the following settings: maximum delta Cn 0.05; decoy database search target False Discovery Rate—FDR 1%; validation based on q-value. Protein-label-free quantitation was performed with the Minora feature detector node at the processing step. Precursor ion quantification used the processing step with the following parameters: Peptides: unique plus razor; precursor abundance was based on intensity; normalization mode was based on the total peptide amount; the pairwise protein ratio calculation and hypothesis test were based on a *t*-test (background based). The Feature Mapper node from the Proteome Discoverer software was used to create features from unique peptide-specific peaks within a short retention time and mass range. This was achieved by applying a chromatographic retention time alignment with a maximum shift of 10 min and 10 ppm of mass tolerance allowed for mapping features from different sample files. For feature linking and mapping, the minimum signal to noise (S/N) threshold was set at 5.

For the determination of DE between groups, the following filters were used: (1) only master proteins detected with high/medium confidence FDR; (2) a protein/phosphoprotein must be detected in more than 50% of samples in each experimental group (except for proteins that were depleted entirely in one of the experimental groups); (3) the *p*-value adjusted using Benjamini–Hochberg correction for the FDR was set to ≤ 0.05; (4) at least 50% of samples with protein-related peptides sequenced by MS/MS; (5) the peptide spectrum matches (PSMs) was set to ≥ 2.

#### Proteomics network analyses

For constructing similarity matrices, DE proteins retrieved from the LFQ experience were uploaded to SynGO (a systematic knowledgebase of functionally-annotated synaptic genes based exclusively on published experimental data (https://syngoportal.org ^69^)) to map Rac1^fl/fl^ and Cx3cr1^CreER+^:Rac1^fl/fl^ synaptosomes (proteins and GO terms). A one graph-based ^70^ pairwise semantic similarities based on the topology of the GO graph structure (Cellular Component ontology) was computed using GoSemSim ^71^ in the R package within Bioconductor ^72^. The same package was used to calculate the semantic similarity among protein clusters using the Best-Match Average (BMA) strategy, aggregating various semantic similarity scores of multiple associated GO terms. Agglomerative Hierarchical clustering using the Ward Linkage was also performed for the similarity matrices.

For network integration and synaptic topographic maps, the set of DE synaptosomal proteins and phosphoproteins were first inputted in SynGO. The DE proteins/phosphoproteins were flagged to presynaptic or postsynaptic compartments using SynGO cellular components’ anthology (dataset release version 20210225). DE proteins of the proteomic dataset positively mapped to SynGO were subsequently screened for synaptic interactors/partners using protein-binding interfaces via inter-protein cross-links in the Stitch database (https://xlink.cncr.nl ^73^). DE phosphoproteins of the phosphoproteomic dataset were inspected and cross-checked using the comprehensive web portal PhosphositePlus (https://www.phosphosite.org/homeAction.action ^74^). Putative protein kinases controlling the phosphorylation of the filtered phosphopeptides were identified using NetworKIN 3.0 (http://networkin.info/index.shtml ^75^). DE phosphoproteins were then screened for synaptic interactors/partners using Stitch and Mechismo ^76^. Pathway enrichment analyses (FDR cutoff < 0.05 with the Benjamini-Hochberg multiple test adjustments) were carried out in WebGestalt ^67^ with REACTOME and GO as functional databases. Network construction and clustering were carried out using k-means clustering in STRING ^68^ and Omics Visualizer ^77^ in Cytoscape.

#### Microglial cell line, live-cell imaging, and FRET

The human microglial cell line HMC3 was obtained from ATCC (ATCC^®^ CRL-3304^™^). Cells were cultivated and maintained as before ^4, 65, 78^. HMC3 microglia were plated on plastic-bottom culture dishes (μ-Dish 35 mm, iBidi) and transfected, as before ^78^, with different biosensors.

Imaging was performed using a Leica DMI6000B inverted microscope. The excitation light source was a mercury metal halide bulb integrated with an EL6000 light attenuator. High-speed, low vibration external filter wheels (equipped with CFP/YFP/farRed excitation and emission filters) were mounted on the microscope (Fast Filter Wheels, Leica Microsystems). A 440-520nm dichroic mirror (CG1, Leica Microsystems) and a PlanApo 63X 1.3NA glycerol immersion objective were used for CFP and FRET images. Images were acquired with 2 × 2 binning using a digital CMOS camera (ORCA-Flash4.0 V2, Hamamatsu Photonics). Quantifications were performed in FIJI, as before ^78^, using the precision FRET (PFRET) data processing software package for ImageJ (https://lvg.virginia.edu/digital-downloads/pfret-data-processing-software). The mean gray intensity values from ratio images were normalized using the deltaF/F0 for statistics. Representative images are shown in intensity-modulated display ^78^. Statistical analyses were conducted using repeated measures two-way ANOVA (individual microglial cells were considered the experimental units; statistical significance was defined as P < 0.05) with the Greenhouse-Geisser correction (to combat sphericity violation) and the Šidák correction for multiple comparisons.

#### Immunofluorescence quantification on cultured microglia

HMC3 microglia were cultivated in glass coverslips and transfected, as before ^78^. Coverslips were fixed with 4% PFA and immunolabeling was performed as before ^4, 23, 65^. Then, coverslips were mounted with Immumount and visualized in a Leica DMI6000B inverted epifluorescence microscope using a PlanApo 63X/1.3NA glycerol immersion objective. Images were acquired using a digital CMOS camera (ORCA-Flash4.0 V2, Hamamatsu Photonics).

Rac1 and Rab27a fluorescence were quantified as before ^4, 23, 65^. GDNF quantifications were performed using the precision FRET (PFRET) data processing software package for ImageJ (https://lvg.virginia.edu/digital-downloads/pfret-data-processing-software). The native 32-bit values from ratio images (GDNF over mCherry or YFP) were used for statistics. Ratio images are shown in intensity-modulated display ^78^. Statistical analyses were conducted using unpaired t-test (single cells were considered the experimental units; statistical significance was defined as P < 0.05).

#### Primary microglial cultures and collection of microglia conditioned media (MCM)

Primary cultures of cortical microglia from neonatal Wistar rats were prepared as before ^23, 66^. Cultures were infected with lentiviruses carrying pLKO-empty or pLKO-Rac1 shRNA as before ^4, 23, 65^. Replication-incopentent lentiviral particles were prepared as before ^4, 23, 79^. After viral removal and medium refeed, microglia were cultured for 5 days. Then, the culture medium (MCM) was collected, centrifuged for debris removal (1200 rpm, 5 min), and frozen at −80 °C until used.

#### Primary cultures of cortical neurons and single-spine live imaging

Brain cortices were dissected from embryonic day 18 Wistar rat embryos and dissociated using trypsin (0.25%, v/v). Neurons were plated at a final density of 1×10^4^ to 5×10^4^ cells per dish and cultured in the presence of a glial feeder layer. Cells were cultured in phenol red-free Neurobasal medium supplemented with B27 supplement (1:50, v/v), 25 mM glutamate, 0.5 mM glutamine, and gentamicin (0.12 mg/ml). To prevent glial overgrowth, neuronal cultures were treated with 5 mM cytosine arabinoside after 3 days in vitro and maintained in a humidified incubator with 5% CO_2_/95% air at 37°C for up to 2 weeks, feeding the cells once per week by replacing one-third of the medium. Cortical neurons were plated on plastic-bottom culture dishes (μ-Dish 35 mm, iBidi) and co-transfected using the calcium phosphate method, as before ^4, 23, 65^, with mCherry-Lifeact-7 and pGP-CMV-GCaMP6f. Experiments were carried out by recording neuronal cultures with microglia conditioned media (MCM) for 10 minutes (baseline) and then in the presence of MCM + KCl 2M (neuronal stimulation).

Imaging was performed using a Leica DMI6000B inverted microscope. The excitation light source was a mercury metal halide bulb integrated with an EL6000 light attenuator. The L5 (Ex: 460-500; BS: 505; Em: 512-542) and TX2 (Ex: 540-580; BS: 595; Em: 607-683) filter set and a PlanApo 63X 1.3NA glycerol immersion objective were used. Images were acquired with 2 × 2 binning using a digital CMOS camera (ORCA-Flash4.0 V2, Hamamatsu Photonics). Quantification of spine area was performed in FIJI, as before ^78^. The mean gray intensity values from spine GCaMP6f and spine actin dynamics (lifeact fluorescence signal) were normalized using the □F/F. Representative GCaMP6f images are shown in intensity-modulated display ^78^.

Data for activity-dependent spine remodeling/calcium dynamics over time were fit using a beta growth and decline model (Y=Ym*(1+ (Te - X)/(Te - Tm))*(X/Te)^(Te/(Te-Tm))) for spine expansion and a plateau followed by one-phase exponential decay (Y= IF(X<X0, Y0, Plateau+(Y0-Plateau)*exp(-K*(X-X0)))) for spine retraction. Spines that collapsed or were eliminated during the evoked neuronal activity were excluded from the analysis. The further additional filters were used for data exclusion: spine expansion < 2.5% spine area; spine retraction < 2% spine area; calcium signal < 25% amplitude increase. Linear regression (with the % spine area as the Y variable) was used to predict how changes in activity-dependent calcium and actin dynamics contributed to spine remodeling. Statistical analyses were conducted using unpaired t-test with single spines measurements as the experimental units (statistical significance was defined as P < 0.05). ANCOVA was used to compare slopes inferred from linear regression analyses.

#### Statistics

Experimenters were blinded to genotypes and housing conditions whenever possible. Statistical tests and their respective significance thresholds are detailed in each subsection of the Experimental Procedures (see above). Descriptive statistics for each dataset are defined in Figure Legends. Statistical analysis and graph construction was performed in GraphPad Prism version 9.0.2 for macOS. Finalized figures were assembled in Adobe Illustrator 2020 (version 24.3).

## Supporting information

SuppFig 1

SuppFig 2

SuppFig 3

SuppFig 4

SuppFig 5

SuppFig 6

## Acknowledgments

Work in the JBR laboratory was financed by FEDER—Fundo Europeu de Desenvolvimento Regional funds through the COMPETE 2020—Operacional Programme for Competitiveness and Internationalisation (POCI), Portugal 2020, and by Portuguese funds through FCT—Fundação para a Ciência e a Tecnologia/Ministério da Ciência, Tecnologia e Ensino Superior in the framework of the project POCI-01-0145-FEDER-031318 (PTDC/MED-NEU/31318/2017).

Work in the TS laboratory was financed by FEDER—Fundo Europeu de Desenvolvimento Regional funds through the COMPETE 2020—Operational Programme for Competitiveness and Internationalisation (POCI), Portugal 2020, and by Portuguese funds through FCT—Fundac□ão para a Cie□ncia e a Tecnologia/Ministério da Cie□ncia, Tecnologia e Ensino Superior in the framework of the project POCI-01-0145-FEDER-030647 (PTDC/ SAU-TOX/30647/2017).

IMP acknowledges funding from European Union’s Seventh Framework Programme for research, technological development and demonstration (Marie Curie Actions) under grant agreement no 600375.

CCP and RS hold employment contracts financed by national funds through FCT—in the context of the program-contract described in paragraphs 4, 5, and 6 of art. 23 of Law no. 57/ 2016, of August 29, as amended by Law no. 57/2017 of July 2019.

The authors acknowledge the support of the following i3S Scientific Platforms: Advanced Light Microscopy (ALM), a member of the national infrastructure PPBI-Portuguese Platform of BioImaging; Animal Facility; Cell culture and Genotyping; Genomics; Proteomics; and Translational Cytometry.

## Supplementary Figure Legends

**Supp. Figure 1**. Microglial Rac1 ablation and relation with known microglial transcriptomic programs (<u>related to Figure 1</u>).

**A**, Strategy for conditionally ablating Rac1 in brain microglia. **B**, Gating strategy for microglial isolation by flow cytometry. **C**, Rac1 mRNA amounts evaluated by qRT-PCR in microglia sorted from the brains of Rac1^fl/fl^ and Cx3cr1^CreER+^:Rac1^fl/fl^ mice (n=3 mice per genotype). Violin plots display medians with quartiles. *p<0.05 (Mann-Whitney test). **D**, Venn diagrams with contingency analyses between the differentially expressed transcripts (RNA-seq) in Cx3cr1^CreER+^:Rac1^fl/fl^ vs. Rac1^fl/fl^ and published microglial transcriptomic programs.

**Supp. Figure 2**. Microglial qRT-PCR transcripts (<u>related to Figure 1</u>).

mRNA amounts evaluated by qRT-PCR in microglia sorted from the brains of Rac1^fl/fl^ and Cx3cr1^CreER+^:Rac1^fl/fl^ mice (n=5-6 mice per genotype). Violin plots display medians with quartiles. *p<0.05; **p<0.01; ***p<0.001 (Mann-Whitney test).

**Supp. Figure 3**. Rac1 controls a Rab27a/GDNF signaling axis in microglia (<u>related to Figure 1</u>).

**A, F and H**, Imaris rendering of confocal images from neocortical tissue sections of Rac1^fl/fl^ and Cx3cr1^CreER+^:Rac1^fl/fl^ mice (3 animals per genotype). Scale bars: 10 μm. **B, D, E, G, I, and J**, Immunofluorescence of Rac1, GDNF, or Rab27a in HMC3 microglia expressing different plasmids (scrambled-mCherry, pLKO-empty, Rac1-shRNA_mCherry, pLKO-Rab27a shRNA, FKBP-YFP+FRBLyn, or Rac1 YF+FRBLyn). Violin plots display medians with quartiles (n=36-58 cells pooled across 3 different cultures). Pseudocolor ramps display representative fluorescence ratio values. **p<0.01; ****p<0.0001 (Mann-Whitney test). Scale bars: 20 μm (B); 10 μm (D, E, G, I, and J). **C**, Flow cytometry analysis of GDNF expression in microglia from Rac1^fl/fl^ and Cx3cr1^CreER+^:Rac1^fl/fl^ mice (n=5 animals per genotype). Violin plots (median and quartiles) depict GDNF^+^ microglia. *P<0.05 (Mann-Whitney test).

**Supp. Figure 4**. Synaptic PPI network controlled by microglial Rac1 (<u>related to Figure 3</u>).

Pre and postsynaptic PPI clusters relative to Cx3cr1^CreER+^:Rac1^fl/fl^ vs. Rac1^fl/fl^ mice. Network displays selected protein nodes (integration of proteome and phosphoproteome datasets [based on Figure 3]) in each cluster.

**Supp. Figure 5**. Microglial Rac1 modulates single spine Ca^2+^ and actin dynamics (<u>related to Figure 3</u>).

**A**, Strategy for studying the impact of microglial Rac1 secretome on spine Ca^2+^ dynamics and actin polymerization/depolymerization. **B-N**, Primary cortical neuronal cultures co-expressing GCamp6f (representative images in C) and Lifeact-DsRed (representative images in D) were recorded in saline for 10 min and then recorded upon stimulation with conditioned media from primary microglial cultures (MCM) carrying scrambled sequence or Rac1 shRNA. In all conditions, neurons were co-stimulated with 2M KCl. Histograms display linear regressions and violin plots display median with quartiles (n=88-112 spines pooled across 3 independent neuronal cultures). ****p<0.0001 (unpaired t test (E and F); ANCOVA (H, I, J, K, L, and M). Scale bars: C: 5 μm; D: 2 μm.

**Supp. Figure 6**. Impact of microglial Rac1 ablation on neuronal numbers, synaptic pruning, and neuroinflammation (<u>related to Figure 3</u>).

**A**, Strategy for preventing aberrant spine Ca^2+^ dynamics elicited by Rac1-deficient microglial secretome. **B**, Primary cortical neuronal cultures expressing GCamp6f were recorded in saline for 10 min and then recorded upon stimulation with conditioned media from Rac1-deficient primary microglial cultures (MCM). In all conditions, neurons were co-stimulated with 20 mM KCl. The MCM was supplemented with GDNF (100 ng/ml) in some conditions. Violin plots (median and quartiles) show single-spine time-lapse fluorescence changes (n=22-34 spines pooled across 3 independent neuronal cultures). *p<0.05; ****p<0.0001 (Mann-Whitney test).**C and D**, Histological confocal analysis of NeuN or Iba1/CD68/PSD-95 on tissue sections from neocortices of Rac1^fl/fl^ and Cx3cr1^CreER+^:Rac1^fl/fl^ mice (n= 3 slices from 4 animals per genotype (C) or n=24 cells from 4 different mice per genotype (D)). **E**, mRNA amounts evaluated by qRT-PCR in neocortices of Rac1^fl/fl^ and Cx3cr1^CreER+^:Rac1^fl/fl^ mice (n=5-6 mice per genotype). Violin plots display medians with quartiles. Scale bars: C: 50 μm; D: 10 μm.

## Notes

### Competing Interest Statement

The authors have declared no competing interest.

