## Supplementary figures and images for "Rac1 signaling in microglia is essential for synaptic proteome plasticity and experience-dependent cognitive performance"

### SuppFig 1

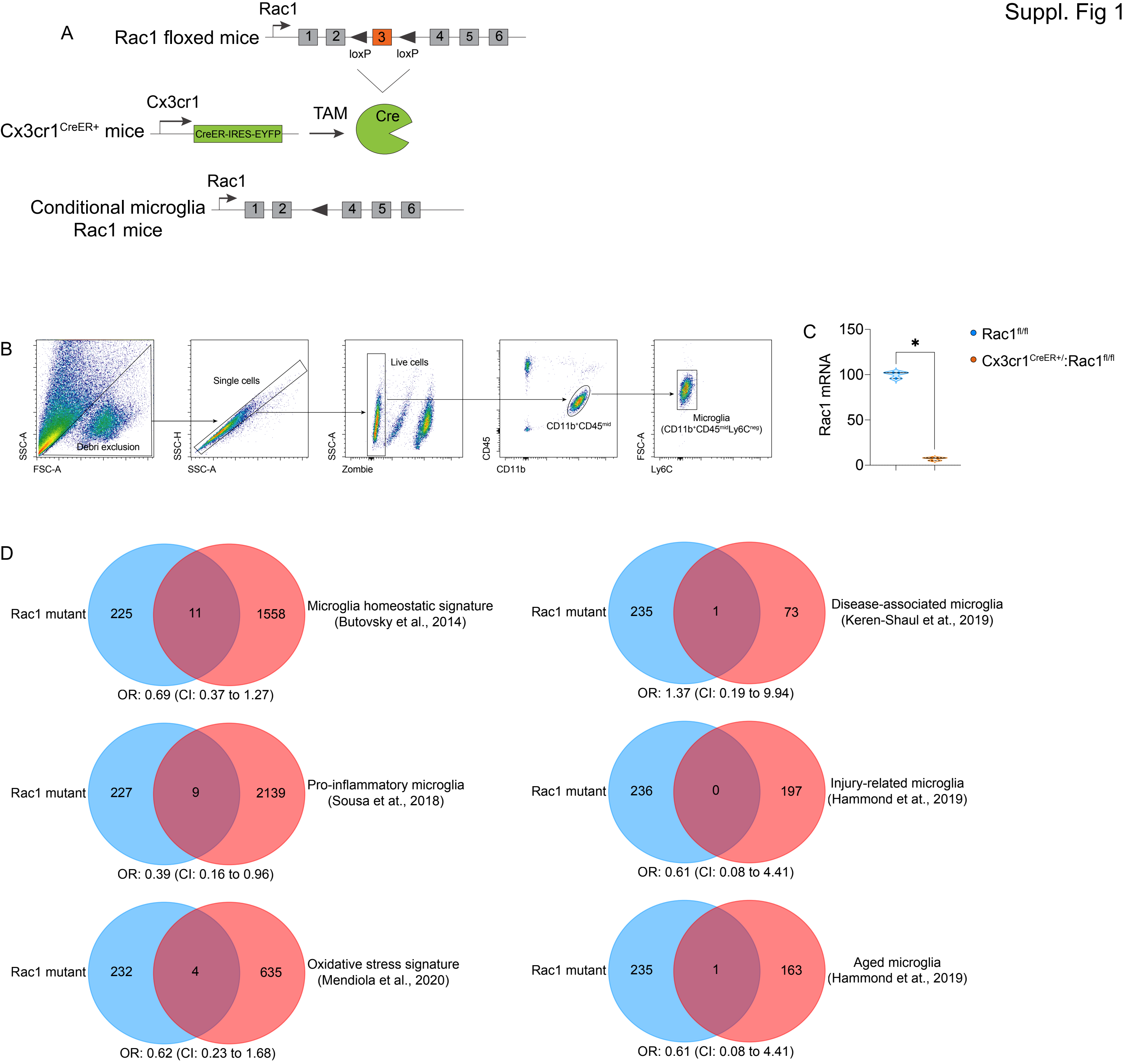

### SuppFig 2

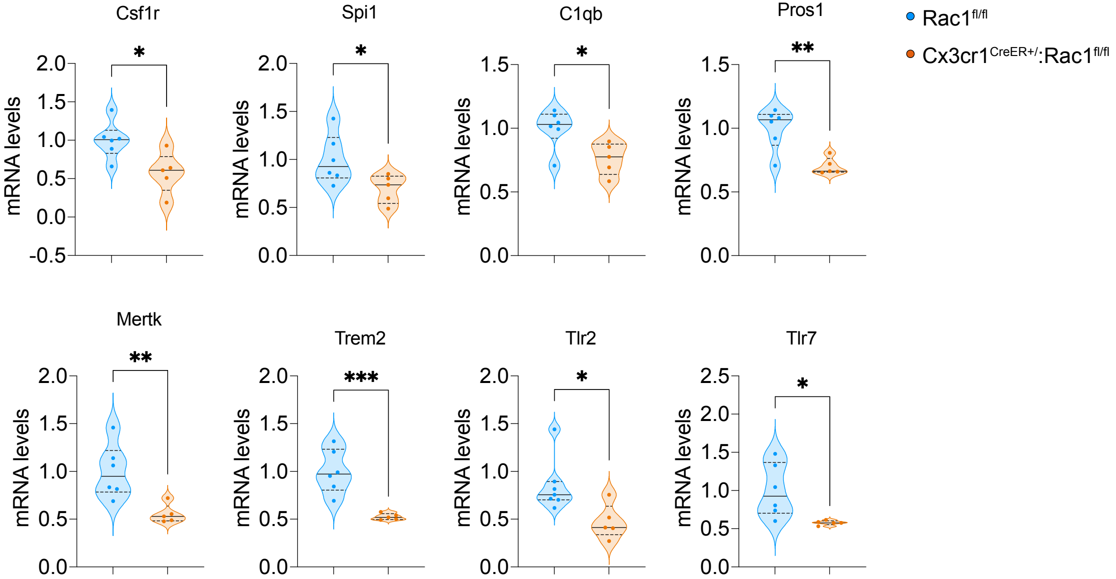

### SuppFig 3

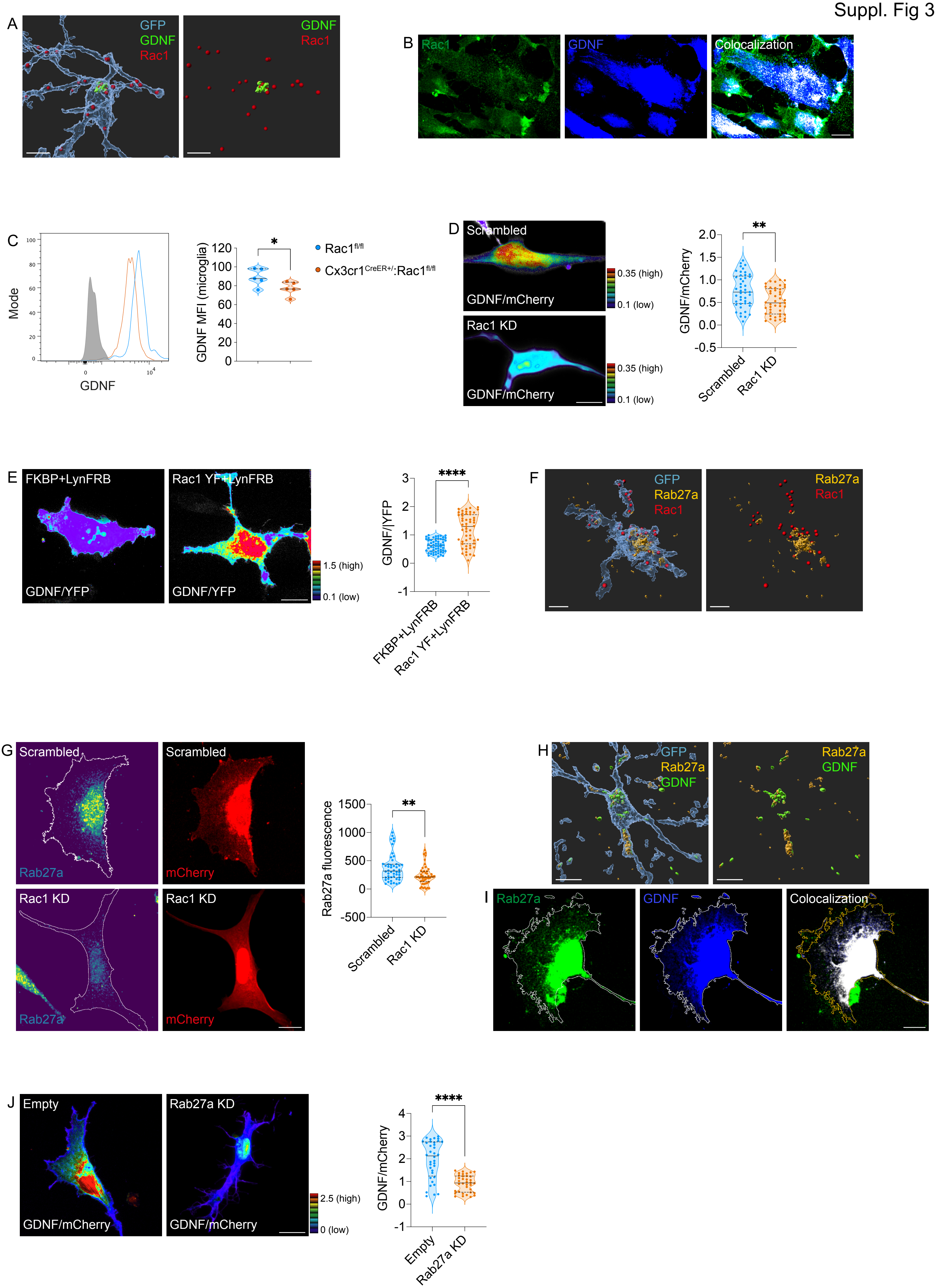

### SuppFig 5

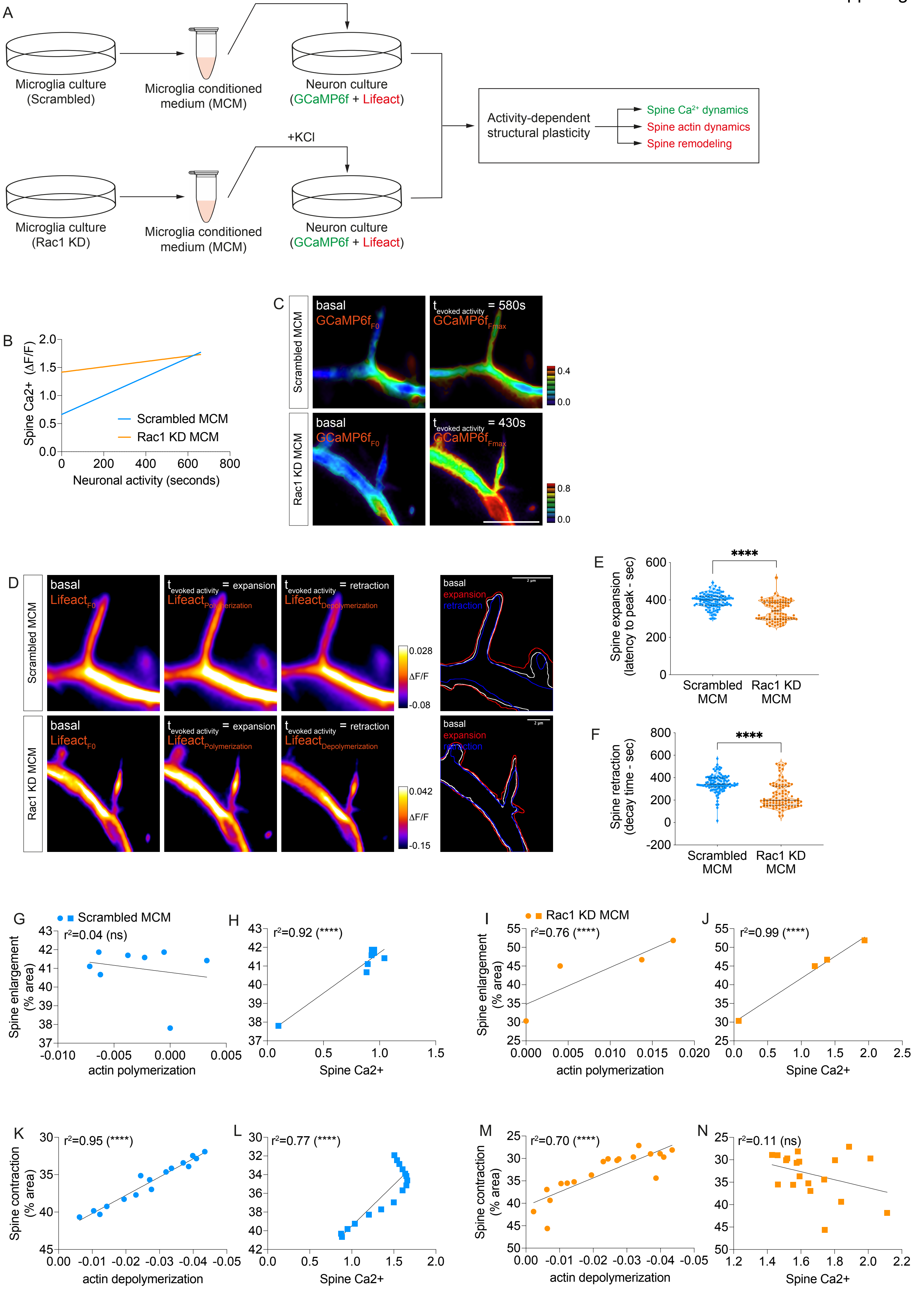

### SuppFig 6

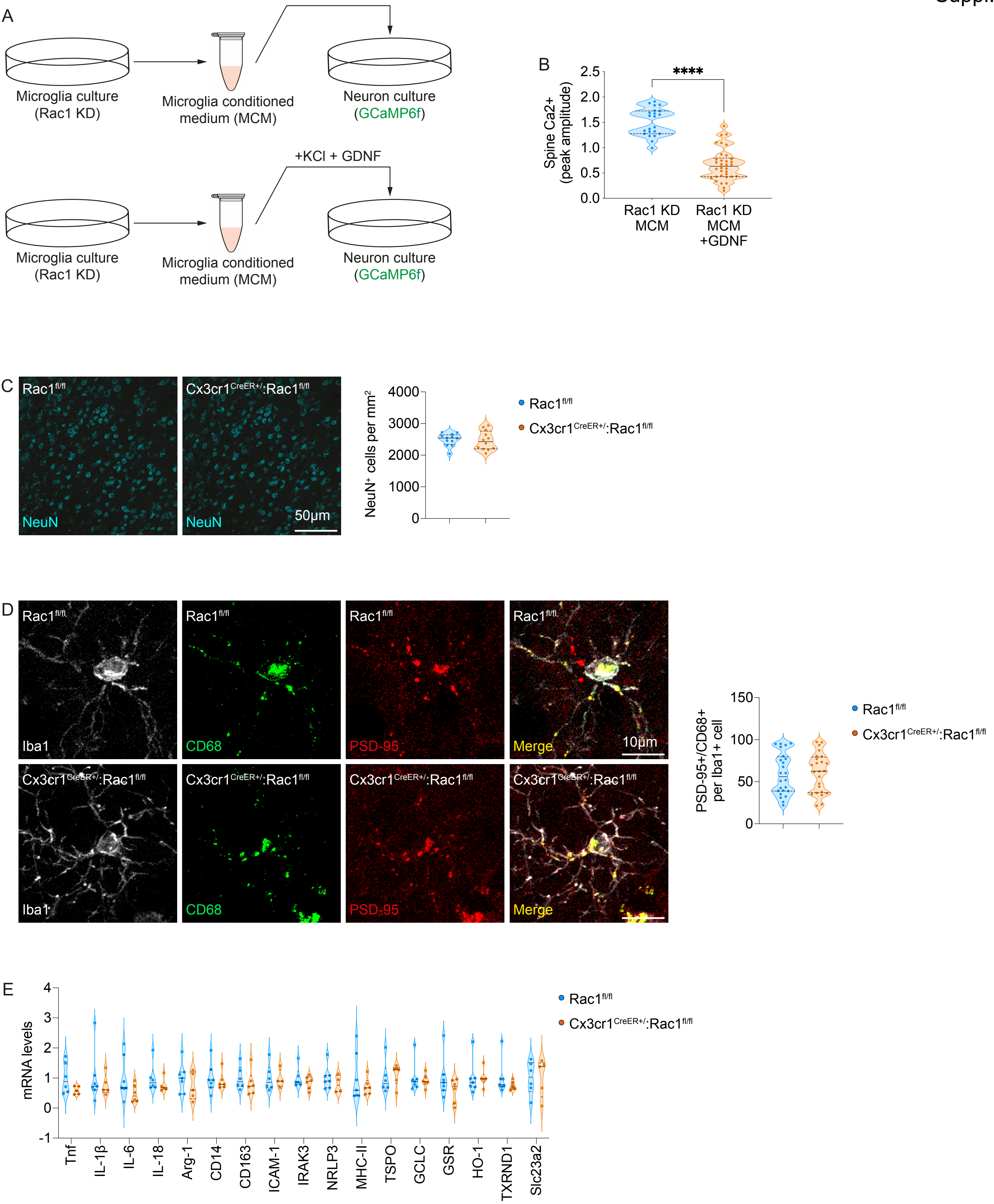
