## Supplementary material for "Rac1 signaling in microglia is essential for synaptic proteome plasticity and experience-dependent cognitive performance": SuppFig 4

Presynaptic network affected by microglial Rac1 signaling

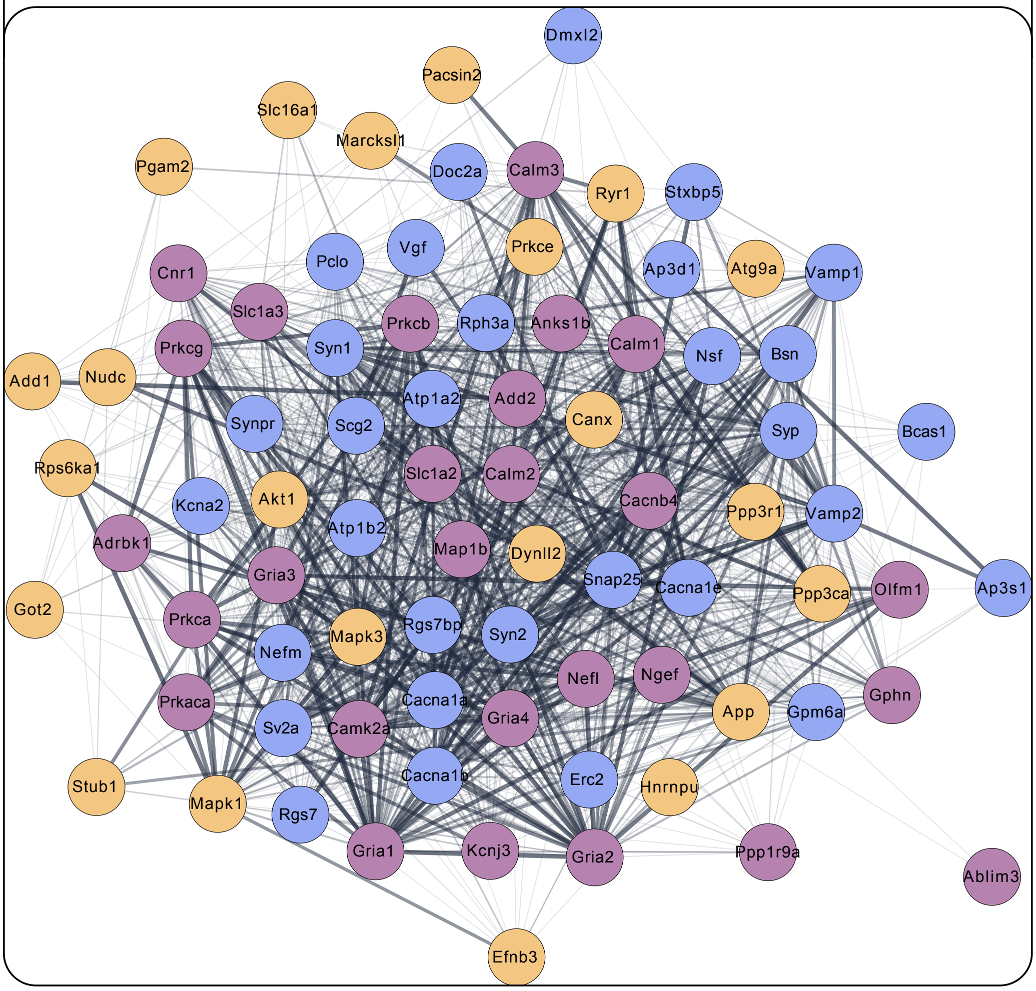


Neurotransmitter release
Presynaptic AMPAr signaling
Lysophospholipid/EPO/IL-1/9

Postsynaptic network affected by microglial Rac1 signaling

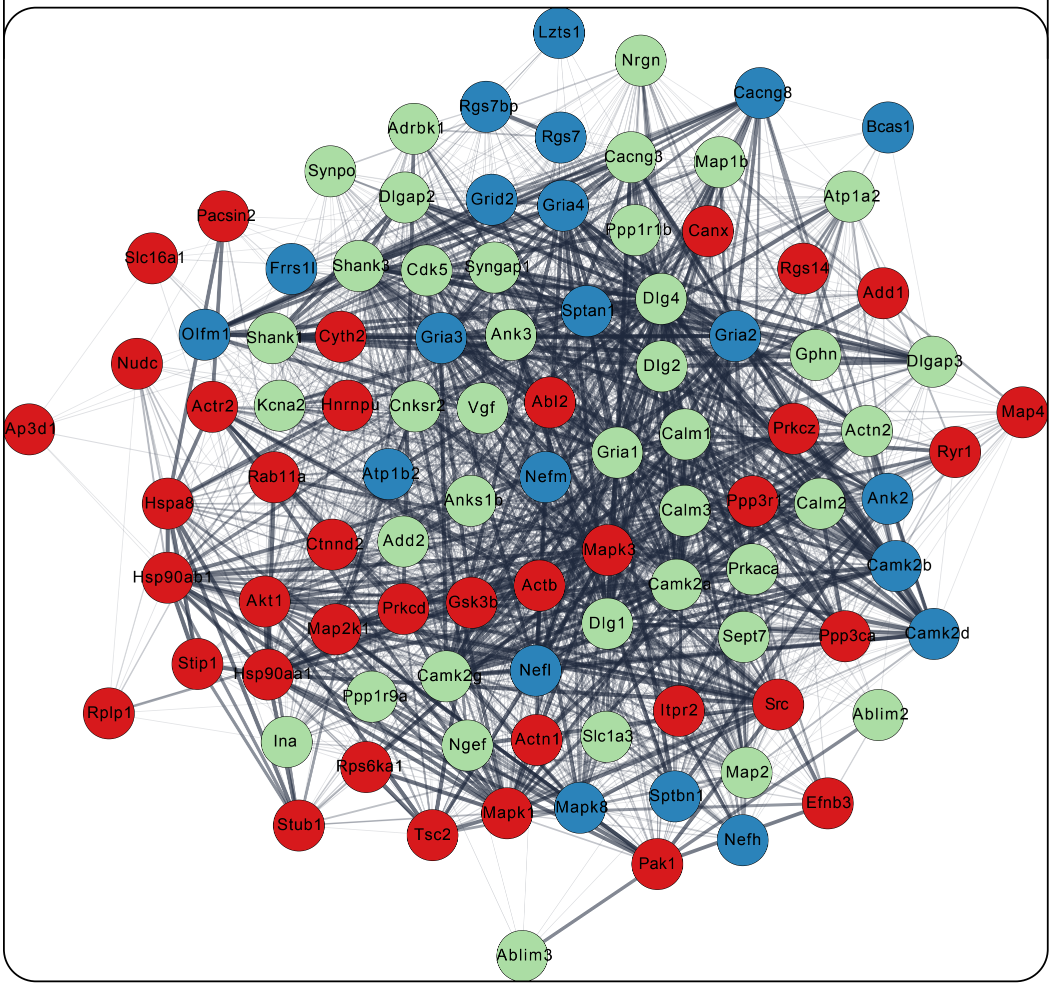


NMDAr-to-CREB signaling
Postsynaptic AMPAr activation
ECM/growth factor signaling
